# The evolution of social learning as phenotypic cue integration

**DOI:** 10.1101/2020.10.30.358028

**Authors:** Bram Kuijper, Olof Leimar, Peter Hammerstein, John M. McNamara, Sasha R. X. Dall

## Abstract

Most analyses of the origins of cultural evolution focus on when and where social learning prevails over individual learning, overlooking the fact that there are other developmental inputs that influence phenotypic fit to the selective environment. This raises the question how the presence of other cue ‘channels’ affects the scope for social learning. Here, we present a model that considers the simultaneous evolution of (i) multiple forms of social learning (involving vertical or horizontal learning based on either prestige or conformity biases) within the broader context of other evolving inputs on phenotype determination, including (ii) heritable epigenetic factors, (iii) individual learning, (iv) environmental and cascading maternal effects, (v) conservative bet-hedging and (vi) genetic cues. In fluctuating environments that are autocorrelated (and hence predictable), we find that social learning from members of the same generation (horizontal social learning) explains the large majority of phenotypic variation, whereas other cues are much less important. Moreover, social learning based on prestige biases typically prevails in positively autocorrelated environments, whereas conformity biases prevail in negatively autocorrelated environments. Only when environments are unpredictable or horizontal social learning is characterised by an intrinsically low information content, other cues such as conservative bet-hedging or vertical prestige biases prevail.

## 1 Introduction

Social learning, the ability to acquire information from other individuals, is a fundamental requirement to cultural evolution [1–4]. Indeed, a substantial amount of theoretical work has sought to identify the ecological conditions in which selection favours social learning as opposed to individual learning (see [3, 4] for reviews), finding that a mixture of both social and individual learning is expected to evolve in fluctuating environments (e.g., [5–9]). However, the conventional focus on the evolution of social versus individual learning overlooks that individuals can also obtain information about their environments by other means. For example, in spatially varying environments an individual’s genotype can become statistically associated to its environment through local adaptation [10, 11], favouring the evolution of genetic cues for phenotypic development [12–15]. In environments that fluctuate predictably over time, theory predicts that individuals are selectively favoured to rely on transgenerational cues that stem from their parent’s phenotype or the parental environment (e.g., [16–20]), transmitted through heritable DNA/histone modifications, parental hormones or even parent-offspring teaching [21–24]. Next, rather than relying on direct or indirect cues about the environment, individuals may also be selectively favoured to rely on mechanisms that generate phenotypic variation instead (e.g., bet-hedging: [25, 26]). Consequently, the availability of cues other than social or individual learning raises the question how organisms should integrate multiple cues when adapting to different environments and how, in turn, this affects the evolutionary scope for social versus individual learning.

While there are a large number of theoretical studies which have analysed the evolutionary implications of subsets of two or three developmental cues (e.g., [12, 27–30]), only a limited number of these studies have sought to predict how organisms should integrate a larger number of available cues [31–33]. These studies find that the rate of environmental change and the degree of environmental predictability are key parameters in determining which cue is most important in phenotype determination, while reliance on mixtures of multiple cues typically occurs in more restrictive settings. However, these studies have only focused on integration of individual and parental cues (e.g., genetic cues, bet-hedging, parental effects and individual learning [phenotypic plasticity]). By contrast, the potential to acquire information via different socially learned cues is yet to be considered in a context that tracks the joint evolution of multiple cues.

To understand how the integration of multiple cues affects the evolution of different social learning mechanisms, we develop a model that tracks the evolution of alternative behavioural phenotypes in a spatiotemporally fluctuating environment. Organisms are selected to develop a behaviour that closely matches the local environment by evolving sensitivity to a range of different cues, where each of these cues potentially provides information about the local environment. Foremost, we consider that individuals can evolve sensitivity to socially learned cues about the phenotypes of others, who are either members of the parental generation (vertical social learning) or of the current generation (horizontal social learning). We then ask how vertical and horizontal social learning evolve jointly with other cues that may affect behavioural development, be it genetic cues, individually learned cues (here represented by within-generational phenotypic plasticity) and transgenerational cues for phenotype determination as in previous models [31–33]. Moreover, because both horizontal and vertical social learning can potentially involve different mechanisms to identify from whom individuals should learn [3, 8], the current model allows sensitivity to evolve based on prestige biases (individuals obtain cues from the most successful individual) and/or conformity biases (individuals obtain cues from the most commonly observed phenotype) for vertical and horizontal social learning independently.

Existing theory on the evolution of learning mechanisms (reviewed in [3, 4, 8]) often stresses the role of different costs in driving the evolution of social and individual learning. For example, social learning is typically thought to result in outdated information relative to individual learning, whereas individual learning is considered to take more effort, resulting in a producer-scrounger game over information (e.g., [5, 8, 34–37]). The key focus of the current model is different, however, as we want to assess what information different combinations of cues can provide and whether some cues can inherently provide more information over and above others. Consequently, we make no a priori assumptions about the relative costs and benefits of one type of cue versus the others, but rather have those payoffs emerge from the ecological scenarios (via migration and timing of life-history events) that impact the information content of the various cues.

## 2 The model

We performed individual-based simulations of a sexually reproducing population distributed over *N*_*p*_ = 40 patches, each supporting a local population of *K* = 100 diploid, hermaphroditic individuals, largely based on a previous model on the evolution of transgenerational effects [17]. While generations are non-overlapping in the sense that only individuals born during the current time step reproduce, the model allows offspring to obtain information from individuals from the previous generation through parental effects and vertical social learning. The simulations are written in `C++` and the code is available at https://doi.org/10.5281/zenodo.3924688. Figure 1A gives an overview of the different cue integration dynamics, while a more elaborate description is provided in section S2 of the online supplement.

**Figure 1.**
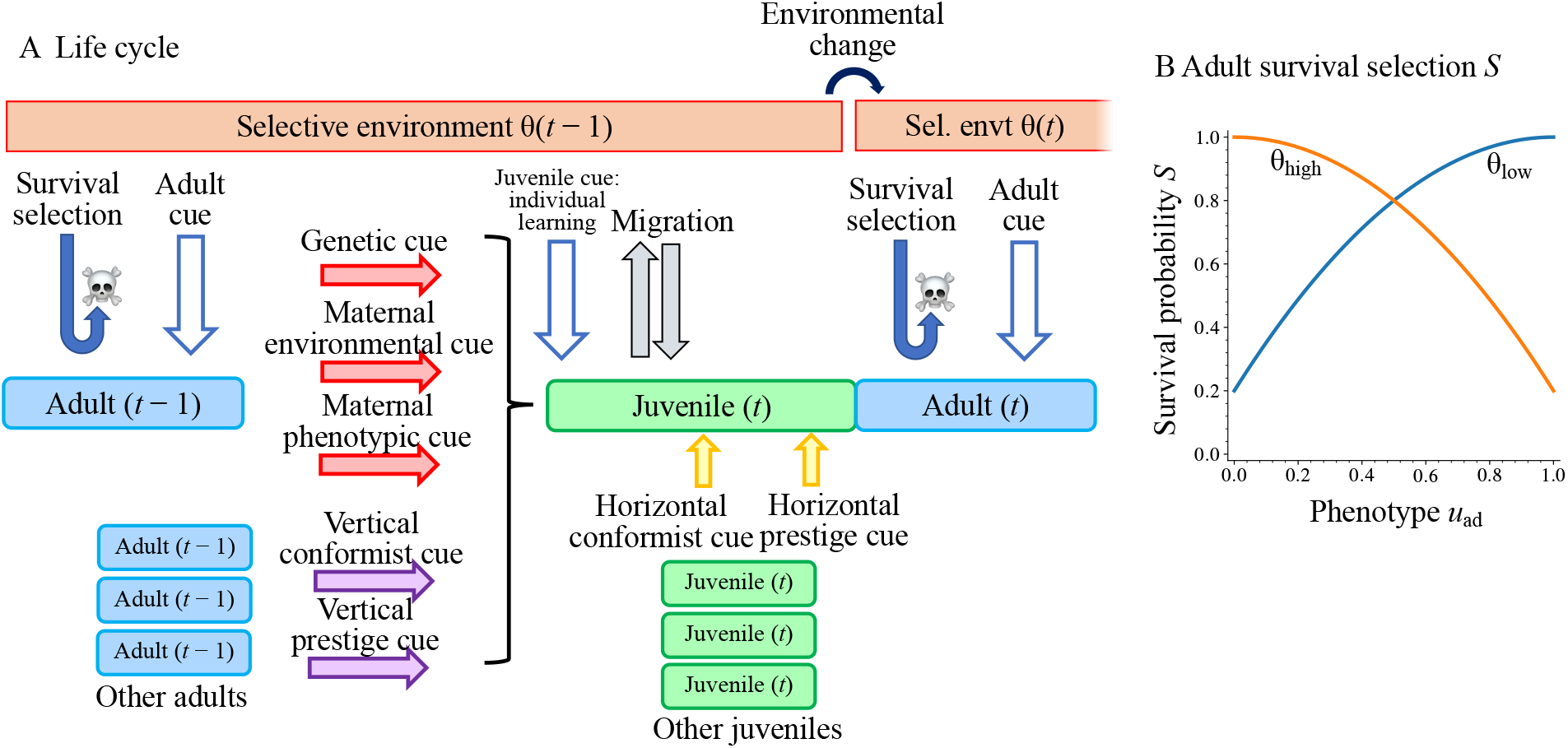
Panel A: The life cycle of the model and the different types of cues: abiotic environmental cues (white arrows), heritable cues (red arrows), vertical social cues (purple arrows) and horizontal social cues (yellow arrows). Panel B: survival probabilities in high and low environments for different adult phenotypes *u*_ad_, where 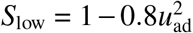 and *S*_high_ = 1 − 0.8 (1 − *u*_ad_)^2^ (following eqns [1,2] in [17]). Throughout the main text, we assume that vertical social learning and individual learning occur prior to migration, while horizontal social learning follows migration. However, we relax these assumptions in the Online Supplement: in Figure S6G-L vertical social learning occurs after migration, in Figure S7A-H horizontal social learning occurs before rather than after migration and in Figure S7G-L individual learning occurs after rather than before migration (see section “future models” in discussion).

### 2.1 Environment

Each patch is either in one of two local environmental states (low: *θ*_low_, or high: *θ*_high_), reflecting, for example, the local temperature or the amount of available resources. Patches can change environmental state independently from other patches at each time step: with probability *p* a patch retains its current environmental state during the next time step, whereas with probability 1 − *p* it changes to the opposite environmental state (similar to two-state models in e.g., [4, 6], but different from models in which the environment continuously varies around an average value [16, 33] or where the environment always attains novel values [4, 6]). Following [17], we assume that both environments change at identical rates, so the global equilibrium frequency *f*_high_ of patches in environmental state *θ*_high_ is given by *f*_high_ = 1 − *f*_low_ = 1/2, while the between-generation environmental autocorrelation of any patch is *ρ*(*θ*_*t*_, *θ*_*t*+1_) = 2*p* − 1, so that when *p* = 0 (rapid change), the autocorrelation is −1, when *p* = 1 (no change), the autocorrelation is 1 and when *p* = 0.5 (random change), the autocorrelation is 0. In Figure S10, we consider values of global equilibrium frequency of patches in environmental state *θ*_high_ other than *f*_high_ = 1/2 (see also [18, 19] for the effect of asymmetries in patch frequencies), but findings are similar to results presented in the main text (e.g., see Figure 2).

**Figure 2.**
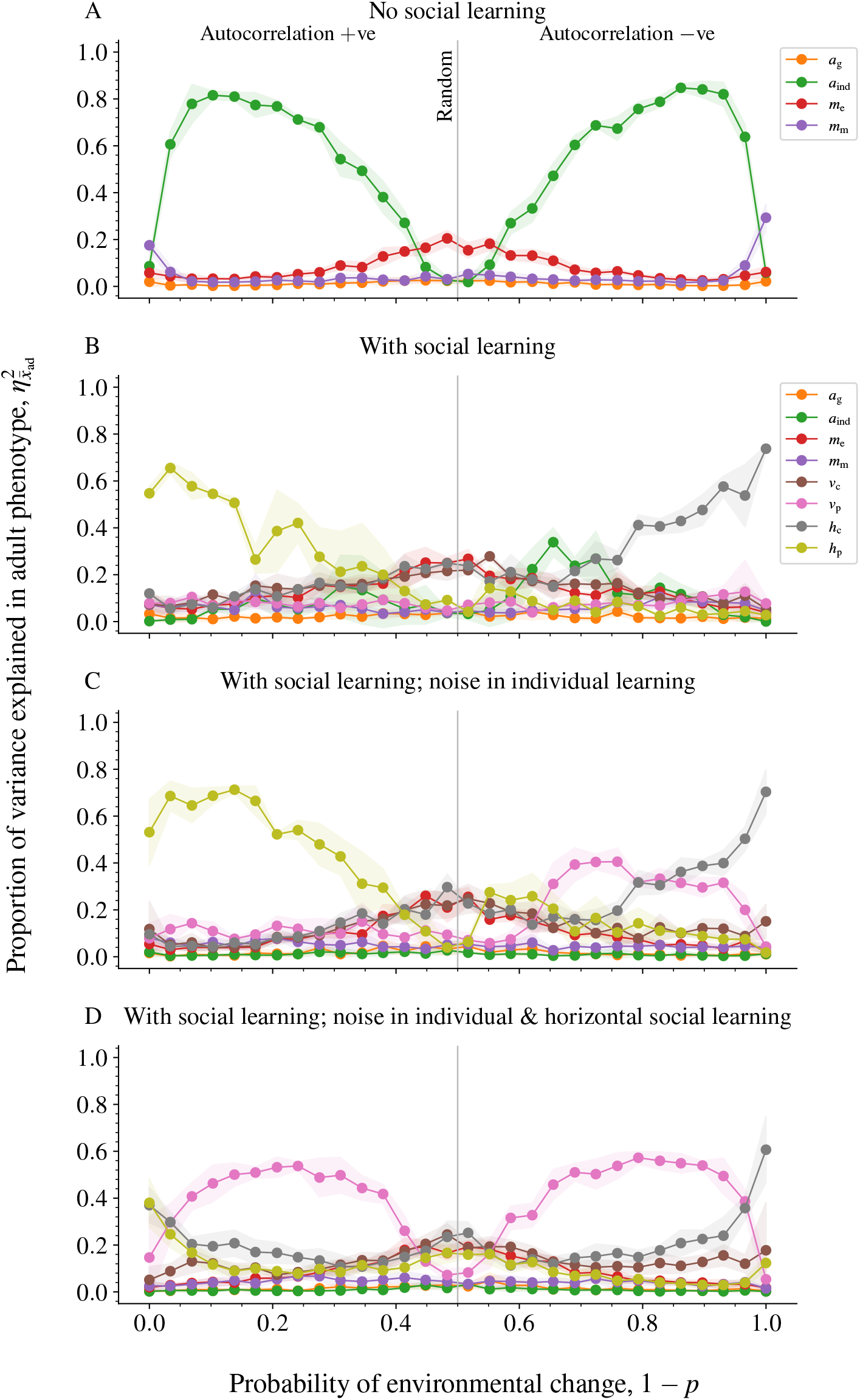
The proportion of variance in adult phenotype (measured at the logistic scale) explained by sensitivity to each cue when varying the probability of environmental change. Each dot reflects the average proportion over *n* = 5 replicate simulations, while envelopes reflect sample standard deviations. Panel A: in case there is substantial noise in horizontal and vertical social learning, individual learning becomes the most important cue on phenotype determination. Panel B: with no noise in horizontally and vertically learned cues, we find that horizontal social learning based on prestige (*h*_*p*_) prevails in positively autocorrelated environments, while individual learning (*a*_ind_) prevails in negatively correlated environments. Interestingly, for strongly negatively autocorrelated environments, we find that a combination of individual learning and horizontal social learning based on conformity (*h*_*c*_) prevails. Panel C: when individually learned environmental cues are un-reliable (*q*_ind_ = 0.5), we find that this has little effect on horizontal social learning based on prestige (*h*_*p*_; compare with panel B), yet individuals now more strongly rely on vertically learned cues based on prestige rather than individually learned cues. Panel D: when both individually learned and horizontally learned cues are unreliable, vertically learned cues based on prestige prevail in autocorrelated environments. Parameters: Panel A: *σ*_h_ = 1.0 (high noise in horizontal social learning), *σ*_*v*_ = 1.0 (high noise in vertical social learning), *q*_ind_ = 1.0 (high fidelity of individual learning). Panel B: *σ*_h_ = *σ*_*v*_ = 0, *q*_ind_ = 1.0. Panel C: *σ*_h_ = *σ*_*v*_ = 0, *q*_ind_ = 0.5. Panel D: *σ*_h_ = 1.0, *σ*_*v*_ = 0, *q*_juv_ = 0.5. Other parameters: *q*_mat_ = 0.5, *d* = 0.1, *n*_*c*_ = *n*_*h*_ = *n* = 5. Variance proportions were calculated through ordinary least squares multiple regression of 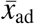 on the right-hand side terms in eq. [3] and then calculating *η*^2^ = SS_between_/SS_total_ for each independent variable (i.e., the partial *R*^2^ also known as the ‘classical’ *η*^2^).

### 2.2 Reproduction and juvenile phenotype determination

Before reproduction, adults experience survival selection based on their adult phenotype *u*_ad_ (see eq. [3] below), where the probability of adult survival *S*(*u*_ad_, *θ*) differs between low and high environments respectively (see Figure 1B and the online supplementary information). Subsequently, *K* newborn offspring are produced by surviving adults. Each newborn offspring is produced by randomly selecting a mother and a father from among the surviving adult breeders, potentially allowing for selfing in case the number of survivors is very small.

Upon birth, an individual offspring then determines its juvenile phenotype *u*_juv_ according to genetic, maternal environmental, maternal phenotypic and vertical social cues (see Figure 1A and eq. 1 below). The juvenile phenotype is also affected by individual learning of the local environment (via juvenile environmental cues), where we assume that individual learning occurs before migration unless indicated otherwise. Consequently, the juvenile phenotype *u*_juv_ that is developed after individual learning is a logistic function of a weighted sum 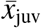 of different cues an individual has received. We have

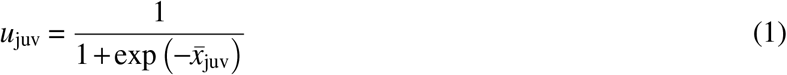

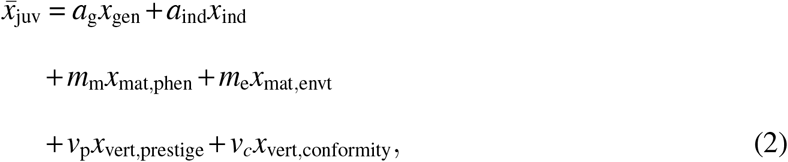

where the *x*_*i*_s are the values of each of the different cues (see section S2.2 in the Online Supplement where we set out the details of the different cues). We then allow the influence of each cue on juvenile phenotype determination to evolve, by assuming that cues are weighed by a set of evolving sensitivity loci, here reflected by variables *a*_g_ (genetic cue), *a*_ind_ (individual learning), *m*_m_ (maternal phenotypic cue), *m*_e_ (maternal environmental cue), *v*_p_ (vertical social learning; prestige bias) and *v*_c_ (vertical social learning; conformity bias) respectively. For the sake of tractability, we assume that each sensitivity locus is diploid and unlinked to other loci. The value of each sensitivity locus is restricted to [−10, 10]. In the absence of any other cues, a sensitivity value *v*_p_ = 0 (vertical social learning sensitivity based on prestige) implies that juveniles attain an intermediate juvenile phenotype of *u*_juv_ = 1/2. A negative value of *v*_p_ implies that when individuals receive a high versus a low value of the *x*_vert,prestige_ cue, they are more likely to develop a low phenotype (*u*_juv_ < 1/2) versus a high phenotype (*u*_juv_ > 1/2). The opposite relationship applies when *v*_p_ is positive.

Regarding the timing and place of the different types of learning, we assume that individual learning occurs in the natal environment prior to migration (see life cycle in Figure 1). However, in Supplementary Figure S7G-L, we consider a scenario in which individual learning occurs after migration to the remote environment. Similar to individual learning, vertical social learning is assumed to occur prior to migration throughout the main text. However, in Supplementary Figure S6G-L, individuals are assumed to perform vertical social learning after migration. By contrast, horizontal social learning is assumed to occur after migration and hence only affects phenotypic development later in life throughout the main text (see eqns. [3,4]). However, in Supplementary Figure S7A-F, we relax these assumptions regarding the timing of horizontal social learning, by assuming that individuals perform horizontal social learning in their natal environment.

After juvenile phenotype determination, individuals migrate with probability *d* to a randomly chosen remote site, while they remain at the natal site with probability 1 − *d*. Consequently, we assume that horizontal social learning (see below) occurs after migration.

### 2.3 Adult phenotype determination

After juvenile phenotype determination, all adults from the previous generation die and individuals learn from other individuals of their current generation (horizontal social learning). As noted before, horizontal social learning occurs only after migration (but see Supplementary Figures S7A - F). Information acquired from horizontally learned social cues is then used by individuals to update the various cue weightings 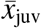 in eq. (2) and develop an adult phenotype. Consequently, the adult phenotype *u*_ad_ that is developed after horizontal social learning is an updated logistic function of a weighted sum 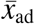 of the various cues an individual has received. We have

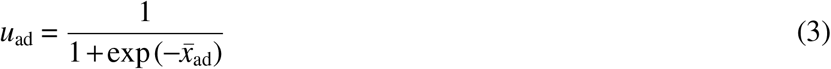

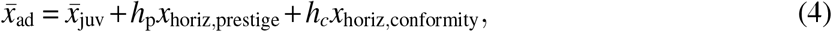

where *h*_p_ (horizontal social learning; prestige bias) and *h*_c_ (horizontal social learning; conformity bias) again reflect unlinked and evolving diploid loci (again bounded between −10 and 10) that reflect sensitivity to both horizontally learned social cues *x*_horiz,prestige_ and *x*_horiz,conformity_ respectively. A full description of the socially learned cues is given in section 2 of the online supplement.

## 3 Results

### 3.1 Result 1: sensitivity to a single cue dominates, but multiple cues are involved in adaptation

Figure 2 depicts four example scenarios -- in which the intrinsic reliability of different cues is varied – that demonstrate how sensitivities to cues jointly evolve in environments that change at different rates of change 1 − *p*. To highlight the relative importance of each cue, Figure 2 shows the proportion of variance in the adult phenotype (measured at the logistic scale) that is explained by sensitivity to each cue, while the evolved values of the sensitivities *a*_*i*_, *m*_*i*_, *h*_*i*_ and *v*_*i*_ (see eqns. [2, 4]) are depicted in Figure S2.

Throughout Figure 2, we find that sensitivity to a single cue explains the large majority of phenotypic variation in adult phenotypes, at least when environments are sufficiently predictable (by being either sufficiently positively or negatively autocorrelated). Moreover, dominant cues are always individually or socially learned, as opposed to genetic cues or maternal effects. In positively autocorrelated environments (left-hand side of each panel in Figure 2), either individual learning (green lines) or social learning driven by prestige biases (horizontal [yellow] or vertical [pink]) prevails. Evolved sensitivities to all other cues explain substantially less phenotypic variance. The prevalence of individual and social learning in predictable environments is unsurprising, as either acquiring direct cues about the environment (individual learning) or obtaining cues about the sampled phenotype with the highest survival in the current environment (prestige-based social learning) provides most information about current conditions.

Only when environments become unpredictable (around the middle of each panel in Figure 2), do individuals start to rely on multiple cues, yet actual sensitivities to the different cues are close to zero (see Figure S2). When environments become largely unpredictable, individuals do not make use of much information at all, but rather develop a conservative bet-hedging strategy with *u*_ad_ = 0.5.

When environments are negatively autocorrelated (right-hand sides of Figure 2B-D), again either individual or social learning prevails. Interestingly, however, if horizontal social learning prevails, it is typically based on conformity biases rather than the prestige biases that are observed in positively autocorrelated environments. Moreover, we find that sometimes multiple cues predominate (e.g., right-hand side of Figure 2C), such as a combination of horizontal social learning based on conformity (grey lines) and vertical social learning (pink lines) based on prestige.

Figures 2B-D show that once the information content of socially learned cues is high enough, they explain the majority of phenotypic variance, while individual learning is far less important. This raises the question what phenotypic information is learned from others in the absence of individual learning, as previous theory suggests that combinations of individual learning and social learning are expected to evolve (e.g., [3, 4, 8]). Figure S3 shows that even when individual learning cannot evolve, social learning still prevails over all other cues. Indeed, social learning still prevails when it only coevolves with genetic cues on phenotype determination (Figure S3A) or maternal cues (Figure S3B). Only when there are no other cues than horizontally learned social cues is there no scope for adaptation (Figure S3C). This is unsurprising, as either individual learning or genes/maternal cues are necessary to result in an adaptive response where phenotypes become associated to their respective environments.

### 3.2 Result 2: timing of environmental change matters

So far, we have considered a scenario where the environment changes between juvenility and adulthood. In Figure S5 we consider a scenario where the environment changes at birth instead, implying that non-migrant juveniles and adults encounter the same environmental conditions. For example, this may reflect fluctuating environments where early-life conditions are highly predictive of the later-life environment [38–40]. We now find that either vertical or horizontal social learning based on prestige biases dominates all other cues for all rates of change (Figure S5B-D), even in a randomly fluctuating environment. As change only happens at birth, the predicted survival of individuals in the current juvenile environment will be highly predictive of adult selective conditions for any rate of change.

Figure 3 generalises the findings above while varying key parameters that affect fidelity of the different cues. Specifically, we vary the fidelity of maternal environmental cues versus individual learning (*x*-axis) and the fidelity of vertical versus horizontal social learning (*y*-axis) for both positively (panels A and C) and negatively autocorrelated environments (panels B and D; see Figure S6 for similar results when varying the migration rate). Similar to Figure S5, Figure 3 shows that horizontal social learning prevails over a large range of the parameter space when the environment changes at birth, as the survival of juveniles is fully informative about the later-life environment (Figure S5C, D). Only when horizontal social learning becomes highly error prone (towards bottom of each panel), is horizontal social learning replaced by vertical social learning. In either case however, social learning is mostly based on prestige biases. Also, we find that there is little difference between positively and negatively autocorrelated environments once the environment changes at birth, as in this case, the rate of change does not affect the relationship between environments experienced between juvenility and adulthood (Figure S5C, D, see also Figure S5).

**Figure 3.**
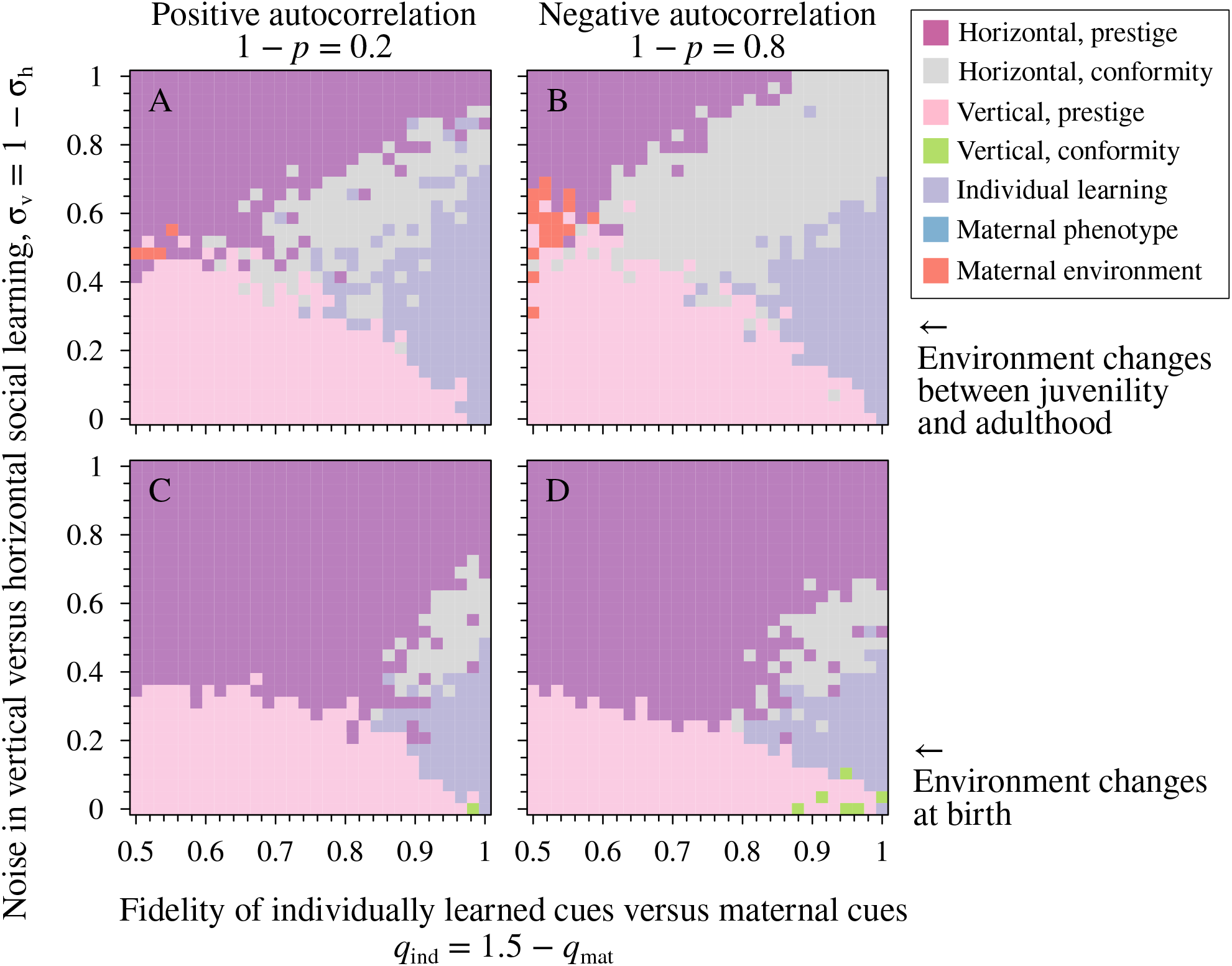
The cue that explains the largest proportion of adult phenotypic variance when varying the fidelity of individually learned versus maternal cues (*x*-axis, with values varying from *q*_ind_ = 0.5*, q*_mat_ = 1.0 [left-hand side] to *q*_ind_ = 1.0*, q*_mat_ = 0.5 [right-hand side]) and the fidelity of vertical versus horizontal socially learned cues (*y*-axis, with values varying from *σ*_vert_ = 0*, σ*_horiz_ = 1 [bottom] to *σ*_vert_ = 1*, σ*_horiz_ = 0 [top]). Panels A, B: environmental change occurs between juvenility and adulthood. Panels C, D: environmental change occurs at birth. Parameters: *d* = 0.1, *n*_c_ = *n*_h_ = 5. Migration rates *d* are varied in Figure S6, while the sample of potential socially learned model individuals *n* = *n*_c_ = *n*_h_ is varied in Figure S8.

By contrast, when the timing of environmental change occurs between juvenility and adulthood, horizontal social learning prevails in a more limited range of parameters (compare Figure 3A, B vs C, D). Interestingly, we find that the fidelity of individual learning affects the evolutionary scope of different social learning mechanisms, as prestige based horizontal social learning predominates when individually learned cues have a low fidelity (*q*_ind_ *≈* 0.5, left hand side of each panel in Figure 3A, B). By contrast, conformity based horizontal social learning predominates once the fidelity of individual learning increases. Moreover, also here we find that conformity is more likely to prevail in negatively autocorrelated environments than in positively autocorrelated ones (see also Figure 2B-D). Finally, in a limited range of parameters where the fidelity of individual learning and horizontal and vertical social learning is low (*σ*_h_ = *σ*_v_ = 0.5, *q*_ind_ = 0.5), we find that environmental maternal effects prevail as it is the only cue that has a considerable fidelity (as *q*_mat_ = 1).

Overall, Figure 3 shows that horizontal social learning (based on either prestige or conformity) often predominates when it comes to the development of the adult phenotype. This raises the question what cues are important in the development of juvenile phenotypes, as those serve as models for horizontal social learning. Figure S4 shows that individual learning and vertical prestige biases (and rarely also maternal effects) are the most important cues in the development of juvenile phenotypes.

## 4 Discussion

Here we provide the first model of how individual and social learning are predicted to coevolve with a multitude of other cues on phenotype determination. Our analysis finds that individual learning or social learning (either horizontal or vertical) typically prevails over all other cues, be it genetic cues, maternal environmental cues, maternal phenotypic cues (i.e., cascading maternal effects) or bet-hedging (i.e., no sensitivity to any cue). Only when cues provide little information about the future do we find that individuals resort to conservative bet-hedging (middle of Figure 2, Supplementary Figure S7C, F). Alternatively, when all other cues prove to be unreliable do we find that environmental maternal effects are selectively favoured (see Figure 3 and Figure S8A).

The prevalence of individual and social learning is to be expected, as individual learning allows individuals to directly detect the state of the local environment. Social learning provides more indirect information about local conditions, as it relies on direct or indirect measures of the performance of others in the local environment: when social learning is based on prestige biases, individuals use cues from the sampled individual with the best performance in the local environment. Figure S8 shows that even when these cues are based on a sample of just *n* = 2 individuals, we still find that socially learned cues (mostly based on conformity biases) prevail in a large range of the parameter space. Being able to rank the performance of others’ phenotypes provides highly accurate information about the local environment, particularly when the sampled phenotypes themselves have accumulated information about the local environment resulting from individual learning or – when individual learning is absent – from selection-based cues that inform about the local environment (genes or cascading maternal effects: see Figure S3).

### 4.1 Social learning in positively versus negatively autocorrelated environments

When environments are negatively autocorrelated (with environmental change occurring between juvenility and adulthood), Figures 2B,C and 3A,B show that conformity biases are considerably more likely to prevail than prestige biases. To understand why horizontal conformity biases prevail over prestige biases, Figure 4 considers the informative value of all learned cues by depicting their correlations with the adult selective environment. Surprisingly, in negatively autocorrelated environments, cues based on horizontal conformism correlate positively (and relatively strongly) with the adult selective environment, whereas horizontal prestige cues are only weakly negatively correlated. Less surprising is that vertically learned and individually learned cues are all negatively correlated with the environment (with individual learned cues exhibiting negative correlations of the strongest magnitude), as the environment experienced by adults of generation *t* − 1 and by juveniles in generation *t* is most likely opposite to the selective environment that will be experienced later by adults in generation *t.*

**Figure 4.**
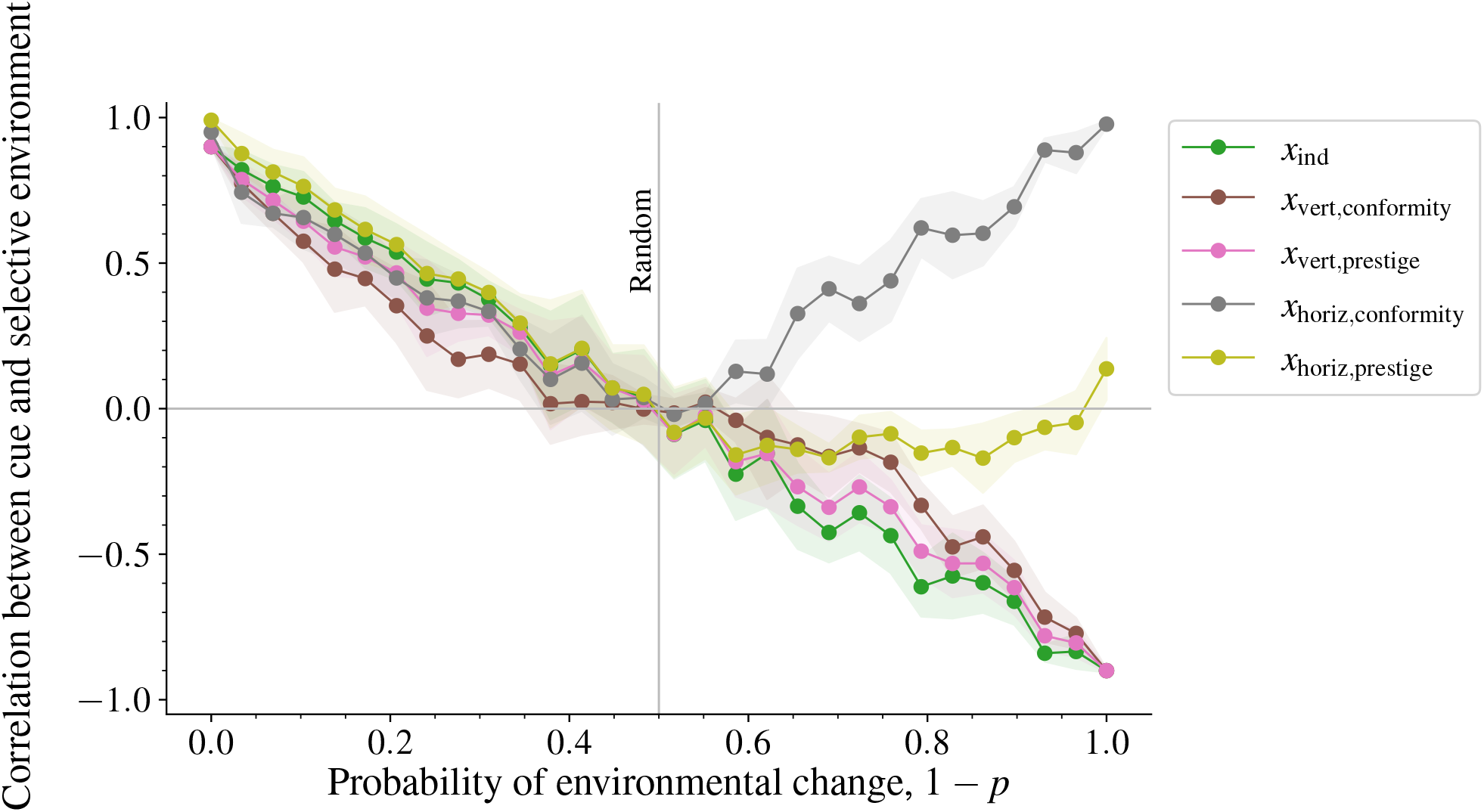
Informational value of the different learned cues. Depicted are the correlations between the adult selective environment *θ*(*t*) and individually and socially learned cues at the preceding juvenile stage. When environments are positively autocorrelated (left-hand side where 1 − *p* < 0.5) we find that vertical conformity-based cues are (on average) the worst predictor of the selective environment, whereas horizontal prestige-based cues are the best predictor of the selective environment. By contrast, in negatively autocorrelated environments, horizontal conformism-based cues are positively correlated with the selective environment, while all other cues are typically negatively correlated. Individually learned cues and conformity-based horizontal social cues have the largest magnitude in negatively autocorrelated environments. Each dot depicts the average correlation over 5 replicate simulations, while envelopes reflect sample standard deviations. Parameters as in Figure 2B.

Why do correlations of these different types of social learning with the selective environment diverge in negatively autocorrelated environments (Figure 4)? Note that juveniles will initially use individual/vertical learning to develop a phenotype opposite to their current juvenile environment. This is because long-term selection (in a negatively autocorrelated environment) has shaped reaction norms to anticipate that selective conditions experienced in adulthood are likely to differ from the environment experienced as a juvenile (indeed, reaction norms based on individual/vertical social learning are negative on the right-hand sides of Figure S2B-D, so that juveniles born in a low environment are likely to develop a phenotype matching a high environment). Once phenotypes based on individual/vertical learning have developed, juveniles perform horizontal social learning. However, as most juveniles now have phenotypes mismatched to their current juvenile environment (but matched to later conditions), individuals are most likely to take prestige-based cues from juvenile models with more intermediate phenotypes (as models with more extreme phenotypes have very low predicted survival values in their juvenile environment and are thus disregarded). In turn, phenotypes of intermediate value are equally likely to occur in any environment, thus resulting in a correlation between horizontal prestige and the adult selective environment of a small magnitude. By contrast, when socially learned cues are based on horizontal conformity biases, a cue is taken that simply reflects the majority of phenotypes without considering any measure of current survival. Consequently, if a majority of sampled juveniles has a low phenotype, this is a good indicator that the adult selective environment will likely be in a low state too. Consequently, conformity-based horizontal social cues become strongly (and positively) correlated with the selective environment.

Figure 3A,B also shows that the fidelity of individually learned cues affects the prevalence of different horizontally learned social cues. When individually learned cues have a low fidelity (*q*_ind_ *≈* 0.5), we find that horizontal prestige based cues predominate, while horizontal conformity based cues prevail otherwise. Why are prestige biased and conformity based social cues differentially affected by the fidelity of individual learning? For low fidelities of individual learning, the distribution of phenotypes in each local deme (before horizontal social learning occurs) is relatively broad, as even modest environmental fluctuations at rates 1 − *p* = 0.2 (Figure 3A) distort strong associations between phenotypes and their local environment created by local adaptation [12, 17]. Consequently, horizontal social learning based on the most frequent phenotype provides little information, as each local deme has a mixture of low and high-adapted phenotypes. By contrast, prestige-based measures of predicted phenotypic performance in the current environment provide a more direct measure of the current environment and therefore prevail. However, once individual learning has a higher fidelity, it allows individuals to modulate their phenotype and match it with the local environment. Consequently, individual learning creates a strong association between the number of individuals with a low versus a high phenotype and their local environment, thus increasing the value of conformity-based horizontal social learning, as this is based on strong differences in the numbers of low versus high phenotypes between both environments.

### 4.2 Why is vertical social learning based on conformity so rare?

Another finding of the current study is that vertical social learning based on conformity rarely predominates, similar to genetic and maternal environmental cues. Vertical conformity biases rarely dominate because they rely on cues about the distribution of parental adult phenotypes, which are, to a large part, a result of cues received in their own juvenile environment at time *t* − 1, resulting in outdated information about adult selection at time *t*. Indeed, Figure 4 shows that the magnitude of the correlation of vertical conformity based cues is small relative to other cues. By contrast, cues based on vertical prestige biases reflect the performance of parental phenotypes in the current (juvenile) environment and are therefore superior to vertical conformism. Moreover, while the relative importance of horizontal prestige versus horizontal conformity is affected by the fidelity of individual learning (see previous paragraph), there is no such interaction between individual learning and the prevalence of vertical prestige vs vertical conformity cues (see bottom of each panel in Figure 3). Again, because individually learned cues provide more recent information about the environment than vertically learned cues, any increase in the fidelity of individual learning tends to replace vertical social learning, rather than affect the evolutionary scope of vertical conformity versus prestige. As found previously [41], the above demonstrates that the order of individual learning versus social learning is likely to strongly affect the evolutionary scope of different forms of social learning.

In some cases, prestige-based cues – which presuppose information of the predicted survival of observed phenotypes – will not be available, so that individuals either have to resort to other means of cue integration. In this case, conformity-based cues may be more general, as they do not rely on direct environmental information, but on a type of ‘crowdsourcing’ in which the most prevalent phenotype of the crowd informs one about the coming environment. Figure S9 shows that when social learning is only based on conformism (while prestige-based cues are excluded), horizontal social learning indeed replaces prestige-based social learning. However, the same does not hold for vertical prestige-based cues, as those are replaced by environmental maternal effects and individual learning. Consequently, we find again that vertical social learning based on conformity does not prevail.

While a direct measure of survival as required by prestige-based cues may typically not be feasible, more indirect measures of prestige are still possible, for example when survival depends on some aspect of phenotypic quality (apart from the phenotype itself; e.g., health or energy reserves) and this quality can be observed, a ranking would be possible. However, the ability to rank others dependent on quality could also imply that a focal individual may have information about its own quality as well, which is something that is not included in the current model. It is likely that personal information about a focal individual’s state may affect the likelihood that it engages in individual or social learning. For example, when genes or maternal effects already provide a good solution for a particular individual, there will not be much reason to copy others. By contrast, when genes or maternal effects provide suboptimal solutions, there will be strong reasons to copy others. Consequently, if cue integration depends on an individual’s state, we would expect strong between-individual differences in cue integration, so that multiple cues are used by the population as a whole. Indeed, such state-dependent [42] social information use may be an important for explaining the existence of consistent individual differences in social learning strategies between individuals [43, 44] and should be a subject of future modelling attempts.

### 4.3 Socially learned cues based on detection versus selection

In a seminal paper by Shea et al. [27] (see also [33]), the information content of different cues has been classified as either selection-based or detection-based, with the aim of explaining differences in inheritance fidelity. Selection-based cues arise when phenotypic variants become correlated to a local environment through differential selection. An example are genetic cues, where local adaptation results in the association of different genetic variants to different selective conditions [12, 13, 15] or maternal phenotypic effects, where selection on the maternal phenotype before reproduction results in associations between the maternal phenotype and the local environment [16, 18, 19, 33]. By contrast, detection-based cues arise when information is directly detected from the environment and subsequently used to modulate phenotypes (which can subsequently be transmitted to offspring). The obvious example of such a detection-based effect is individual learning, but also maternal environmental effects resemble a scenario where a phenotype is only transmitted to offspring once it has been detected [27]. In case of social learning, prestige-based cues are clearly detection-based, as they involve a measure of phenotypic performance in its current environment. By contrast, conformity-based cues can be both selection and detection-based, as conformity is a function of the distribution of the different phenotypes in the local deme: this distribution is both a result of differential survival selection and of individuals modulating their phenotypes via detection-based mechanisms (i.e., individual learning). Consequently, conformity-based social learning can result in either short-term inheritance of variants (when those variants are newly generated each generation through detection-based mechanisms), or it can result in long-term inheritance (when a variant is copied because it is the most frequent due to differential selection).

### 4.4 Future work

The current study has only scratched the surface when it comes to integrating social learning and cultural evolution with other forms of cue integration. To study the interplay of different cue integration mechanisms, our model considered a scenario of local adaptation to a spatiotemporally environment that fluctuates between two types, as has been the subject of numerous previous models of cue integration (e.g.,[17, 20, 32]). However, studies in the context of social learning have considered different environmental configurations, such as a scenario in which the environment changes into a previously unknown state (i.e., similar to ‘infinite states’ models of social learning [4, 6]). It would be interesting to assess the consequences of more continuous forms of environmental variation for the evolution of cue integration. Based on somewhat similar models that considered the effect of large environmental changes to adaptation, we expect that combinations of multiple cues would prevail in such circumstances ([45], Figure 2 in [16]). In particular, we would predict that not only individual learning prevails (to acquire information about the most recent state of the environment; e.g., [4, 6]), but also mechanisms to ensure recently acquired phenotypes are inherited across generations (e.g., (vertical) social learning and cascading maternal effects), as any genetic inputs to the trait are likely to slowly evolve to an ever changing environment and hence be largely outdated. Future studies are needed to consider the evolution of cue integration mechanisms in such environmental configurations.

Also, the timing at which individuals obtain information from different types of cues could substantially affect the outcome (e.g., [36]). For the sake of tractability, our study focused on a scenario in which individuals perform vertical social learning and individual learning before migration, while horizontal social learning was performed following migration (see Figure 1). In Supplementary Figures S6 and S7 we have, however, relaxed these assumptions, for different probabilities of migration. Unsurprisingly, if migration is relatively low, the timing at which learning occurs has little impact and outcomes are very similar to those in Figure 3. When dispersal probabilities are higher, we find that the latest cues received (i.e., those received subsequent to migration) prevail over all others. For example, in Supplementary Figure S6G-L, both vertical and horizontal social learning occur subsequent to migration, whereas individual learning occurs prior to migration. As a consequence, we find that individual learning is nearly absent when migration rates are high (Figure S6I, L). By contrast, when individual learning occurs subsequent to migration, it prevails for a much wider range of the parameter space, particularly when migration rates are high (Figure S7I, L). Finally, when neither cue is received subsequent to migration, we find that conservative bet-hedging prevails when migration rates are high (Figure S7C, F). Hence, as noted in the main results in which we varied the timing of environmental change, the timing of the different learning events also matters for the predominance of different cues over others. We advocate for more studies that systematically vary the timing at which (combinations of) different cues are obtained during the life cycle.

In relation to the timing of cues, our model also assumes that selection only acts during adulthood, prior to reproduction (following previous models: [12, 16, 17]). By contrast, how individuals integrate suits of different cues when selection acts on juveniles vs adults has yet to be assessed. We would expect that selection on juveniles (e.g., prior to horizontal learning) would have a similar effect to setting the timing of environmental change to birth (see Figure S5 and Figure 3C, D). Once selection acts during early life, only those juvenile phenotypes that match the local environment will survive. Consequently, the juvenile phenotype becomes more informative about the later-life environment, thus favouring higher levels of horizontal social learning as in Figure 3C, D. Overall, future studies are needed to systematically analyse how adaptive cue integration varies across different stages of the life cycle.

## Supporting information

Supplementary figures and model description

## Acknowledgements

We would like to thank Piet van den Berg and two anonymous reviewers for suggesting several improvements to this manuscript. BK acknowledges support from the Leverhulme Trust (Research Project Grant RPG 2018-380). The authors would like to acknowledge the use of the University of Exeter’s Advanced Research Computing (ARC-Cornwall) facilities in carrying out this work. OL acknowledges support from the Swedish Research Council (grant number 2018-03772).

## Notes

https://doi.org/10.5281/zenodo.3924688

### Competing Interest Statement

The authors have declared no competing interest.

## References

[1] Cavalli-Sforza, L. L. & Feldman, M. W., 1981 Cultural Transmission and Evolution. Princeton: Princeton University Press. doi:10.2307/j.ctvx5wbt8.

[2] Boyd, R. & Richerson, P. J., 1985 Culture and the Evolutionary Process. Chicago: Chicago University Press.

[3] Henrich, J. & McElreath, R., 2003 The evolution of cultural evolution. Evol. Anthropol. 12, 123–135. doi:10.1002/evan.10110.

[4] Aoki, K. & Feldman, M. W., 2014 Evolution of learning strategies in temporally and spatially variable environments: a review of theory. Theor. Popul. Biol. 91, 3–19. doi:10.1016/j.tpb.2013.10.004.

[5] Rogers, A. R., 1988 Does biology constrain culture? Am. Anthropol. 90, 819–831. doi:10.1525/aa.1988.90.4.02a00030.

[6] Feldman, M. W., Aoki, K. & Kumm, J., 1996 Individual versus social learning: evolutionary analysis in a fluctuating environment. Anthropol. Sci. 104, 209–232. doi:10.1537/ase.104.209.

[7] Aoki, K., Wakano, J. Y. & Feldman, M. W., 2005 The emergence of social learning in a temporally changing environment: a theoretical model. Curr. Anthropol. 46, 334–340. doi:10.1086/428791.

[8] Laland, K. N., 2004 Social learning strategies. Learn. Behav. 32, 4–14. doi:10.3758/bf03196002.

[9] Rendell, L., Fogarty, L. & Laland, K. N., 2010 Roger’s paradox recast and resolved: population structure and the evolution of social learning strategies. Evolution 64, 534–548. doi:10.1111/j.1558-5646.2009.00817.x.

[10] Kawecki, T. J. & Ebert, D., 2004 Conceptual issues in local adaptation. Ecol. Lett. 7, 1225–1241. doi:10.1111/j.1461-0248.2004.00684.x.

[11] Savolainen, O., Lascoux, M. & Merilä, J., 2013 Ecological genomics of local adaptation. Nat. Rev. Genet. 14, 807–820. doi:10.1038/nrg3522.

[12] Leimar, O., Hammerstein, P. & Van Dooren, T. J. M., 2006 A new perspective on developmental plasticity and the principles of adaptive morph determination. Am. Nat. 167, 367–376. doi:10.1086/499566.

[13] Dall, S. R. X., McNamara, J. M. & Leimar, O., 2015 Genes as cues: phenotypic integration of genetic and epigenetic information from a Darwinian perspective. Trends Ecol. Evol. 30, 327–333. doi:10.1016/j.tree.2015.04.002.

[14] Leimar, O., Dall, S. R. X., Hammerstein, P. & McNamara, J. M., 2016 Genes as cues of relatedness and social evolution in heterogeneous environments. PLoS Comput. Biol. 12, e1005 006. doi:10.1371/journal.pcbi.1005006.

[15] Leimar, O., Dall, S. R. X., McNamara, J. M., Kuijper, B. & Hammerstein, P., 2019 Ecological genetic conflict: genetic architecture can shift the balance between local adaptation and plasticity. Am. Nat. 193, 70–80. doi:10.1086/700719.

[16] Kuijper, B. & Hoyle, R. B., 2015 When to rely on maternal effects and when on phenotypic plasticity? Evolution 69, 950–968. doi:10.1111/evo.12635..

[17] Leimar, O. & McNamara, J. M., 2015 The evolution of transgenerational integration of information in heterogeneous environments. Am. Nat. 185, E55–E69. doi:10.1086/679575.

[18] English, S., Pen, I., Shea, N. & Uller, T., 2015 The information value of nongenetic inheritance in plants and animals. PLoS ONE 10, e0116 996. doi:10.1371/journal.pone.0116996.

[19] Kuijper, B. & Johnstone, R. A., 2016 Parental effects and the evolution of phenotypic memory. J. Evol. Biol. 29, 265–276. doi:10.1111/jeb.12778.

[20] Proulx, S. R. & Teotónio, H., 2017 What kind of maternal effects can be selected for in fluctuating environments? Am. Nat. 189, E118–E137. doi:10.1086/691423.

[21] Danchin, É., Charmantier, A., Champagne, F. A., Mesoudi, A., Pujol, B. & Blanchet, S., 2011 Beyond DNA: integrating inclusive inheritance into an extended theory of evolution. Nat. Rev. Genet. 12, 475–486. doi:10.1038/nrg3028.

[22] Radford, E. J., 2018 Exploring the extent and scope of epigenetic inheritance. Nat. Rev. Endocrinol. 14, 345–355. doi:10.1038/s41574-018-0005-5.

[23] Bošković, A. & Rando, O. J., 2018 Transgenerational epigenetic inheritance. Annu. Rev. Genet. 52, 21–41. doi:10.1146/annurev-genet-120417-031404.

[24] Groothuis, T. G. G., Hsu, B.-Y., Kumar, N. & Tschirren, B., 2019 Revisiting mechanisms and functions of prenatal hormone-mediated maternal effects using avian species as a model. Philos. Trans. R. Soc. Lond. B Biol. Sci. 374, 20180 115. doi:10.1098/rstb.2018.0115.

[25] Seger, J. & Brockmann, J. H., 1989 What is bet-hedging? Oxf. Surv. Evol. Biol. 4, 182–211.

[26] Starrfelt, J. & Kokko, H., 2012 Bet-hedging – a triple trade-off between means, variances and correlations. Biol. Rev. 87, 742–755. doi:10.1111/j.1469-185X.2012.00225.x.

[27] Shea, N., Pen, I. & Uller, T., 2011 Three epigenetic information channels and their different roles in evolution. J. Evol. Biol. 24, 1178–1187. doi:10.1111/j.1420-9101.2011.02235.x.

[28] Kuijper, B., Johnstone, R. A. & Townley, S., 2014 The evolution of multivariate maternal effects. PLoS Comput. Biol. 10, e1003 550. doi:10.1371/journal.pcbi.1003550.

[29] Tufto, J., 2015 Genetic evolution, plasticity and bet-hedging as adaptive responses to temporally autocorrelated fluctuating selection: a quantitative genetic model. Evolution 69, 2034–2049. doi:10.1111/evo.12716.

[30] van Gestel, J. & Weissing, F. J., 2016 Regulatory mechanisms link phenotypic plasticity to evolvability. Sci. Rep. 6, 24 524. doi:10.1038/srep24524.

[31] Rivoire, O. & Leibler, S., 2014 A model for the generation and transmission of variations in evolution. Proc. Natl. Acad. Sci. U.S.A. 111, E1940–E1949. doi:10.1073/pnas.1323901111.

[32] Botero, C. A., Weissing, F. J., Wright, J. & Rubenstein, D. R., 2015 Evolutionary tipping points in the capacity to adapt to environmental change. Proc. Natl. Acad. Sci. U.S.A. 112, 184–189. doi:10.1073/pnas.1408589111.

[33] McNamara, J. M., Dall, S. R. X., Hammerstein, P. & Leimar, O., 2016 Detection vs. selection: integration of genetic, epigenetic and environmental cues in fluctuating environments. Ecol. Lett. 19, 1267–1276. doi:10.1111/ele.12663.

[34] Barnard, C. & Sibly, R., 1981 Producers and scroungers: A general model and its application to captive flocks of house sparrows. Anim. Behav. 29, 543–550. doi:10.1016/s0003-3472(81)80117-0.

[35] Giraldeau, L., Valone, T. J. & Templeton, J. J., 2002 Potential disadvantages of using socially acquired information. Philos. Trans. R. Soc. Lond. B Biol. Sci. 357, 1559–1566. doi:10.1098/rstb.2002.1065.

[36] Enquist, M., Eriksson, K. & Ghirlanda, S., 2007 Critical social learning: a solution to Rogers’s paradox of nonadaptive culture. Am. Anthropol. 109, 727–734. doi:10.1525/aa.2007.109.4.727.

[37] Bossan, B., Jann, O. & Hammerstein, P., 2015 The evolution of social learning and its economic consequences. J. Econ. Behav. Organ. 112, 266–288. doi:10.1016/j.jebo.2015.01.010.

[38] Frankenhuis, W. E., Nettle, D. & McNamara, J. M., 2018 Echoes of early life: recent insights from mathematical modeling. Child Dev. 89, 1504–1518. doi:10.1111/cdev.13108.

[39] Kuijper, B., Hanson, M. A., Vitikainen, E. I. K., Marshall, H. H., Ozanne, S. E. & Cant, M. A., 2019 Developing differences: early-life effects and evolutionary medicine. Philos. Trans. R. Soc. Lond. B Biol. Sci. 374, 20190 039. doi:10.1098/rstb.2019.0039.

[40] Gluckman, P. D., Hanson, M. A. & Low, F. M., 2019 Evolutionary and developmental mismatches are consequences of adaptive developmental plasticity in humans and have implications for later disease risk. Philos. Trans. R. Soc. Lond. B Biol. Sci. 374, 20180 109. doi:10.1098/rstb.2018.0109.

[41] Eriksson, K., Enquist, M. & Ghirlanda, S., 2007 Critical points in current theory of conformist social learning. J. Evol. Psych. 5, 67–87. doi:10.1556/jep.2007.1009.

[42] Houston, A. I. & McNamara, J. M., 1999 Models of Adaptive Behaviour: An Approach Based on State. Cambridge: Cambridge University Press.

[43] Molleman, L., van den Berg, P. & Weissing, F. J., 2014 Consistent individual differences in human social learning strategies. Nat. Commun. 5. doi:10.1038/ncomms4570.

[44] Mesoudi, A., Chang, L., Dall, S. R. X. & Thornton, A., 2016 The evolution of individual and cultural variation in social learning. Trends Ecol. Evol. 31, 215–225. doi:10.1016/j.tree.2015.12.012.

[45] Lande, R., 2009 Adaptation to an extraordinary environment by evolution of phenotypic plasticity and genetic assimilation. J. Evol. Biol. 22, 1435–1446. doi:10.1111/j.1420-9101.2009.01754.x..

[46] Kirkpatrick, M. & Lande, R., 1989 The evolution of maternal characters. Evolution 43, 485–503. doi:10.2307/2409054.

[47] McGlothlin, J. W. & Galloway, L. F., 2013 The contribution of maternal effects to selection response: an empirical test of competing models. Evolution 68, 549–558. doi:10.1111/evo.12235.

[48] Rossiter, M. C., 1998 The role of environmental variation in parental effects expression. In Maternal Effects as Adaptations (eds. T. A. Mousseau & C. W. Fox). Oxford: Oxford University Press, pp. 112–134.

[49] Henrich, J., 2004 Demography and cultural evolution: how adaptive cultural processes can produce maladaptive losses—the Tasmanian case. Am. Antiq. 69, 197–214. doi:10.2307/4128416.

[50] Kobayashi, Y. & Aoki, K., 2012 Innovativeness, population size and cumulative cultural evolution. Theor. Popul. Biol. 82, 38–47. doi:10.1016/j.tpb.2012.04.001.

