## Supplementary figures and model description for "The evolution of social learning as phenotypic cue integration"

#### S1 Supplementary Figures

**Figure S1** Example simulations in a negatively autocorrelated environment ($1-p=0.9$), where each line depicts the average trait value of a single replicate run for over 75000 generations. Sensitivity to individually learned cues $a_{\text{ind}}$ evolves to slight negative values (panel B), while both vertical and horizontally learned cues based on conformism ($v_{\text{c}}$ and $h_{\text{c}}$) evolve to positive values (panels D,E). See also panel B of Figure S2 at the value $1-p=0.9$. Parameters: $q_{\text{ind}}=1.0$, $q_{\text{mat}}=0.5$, $n=5$, $m=0.1$, $1-p=0.9$, $\sigma_{\text{h}}=\sigma_{v}=0.0$. Number of replicate simulations: 5.

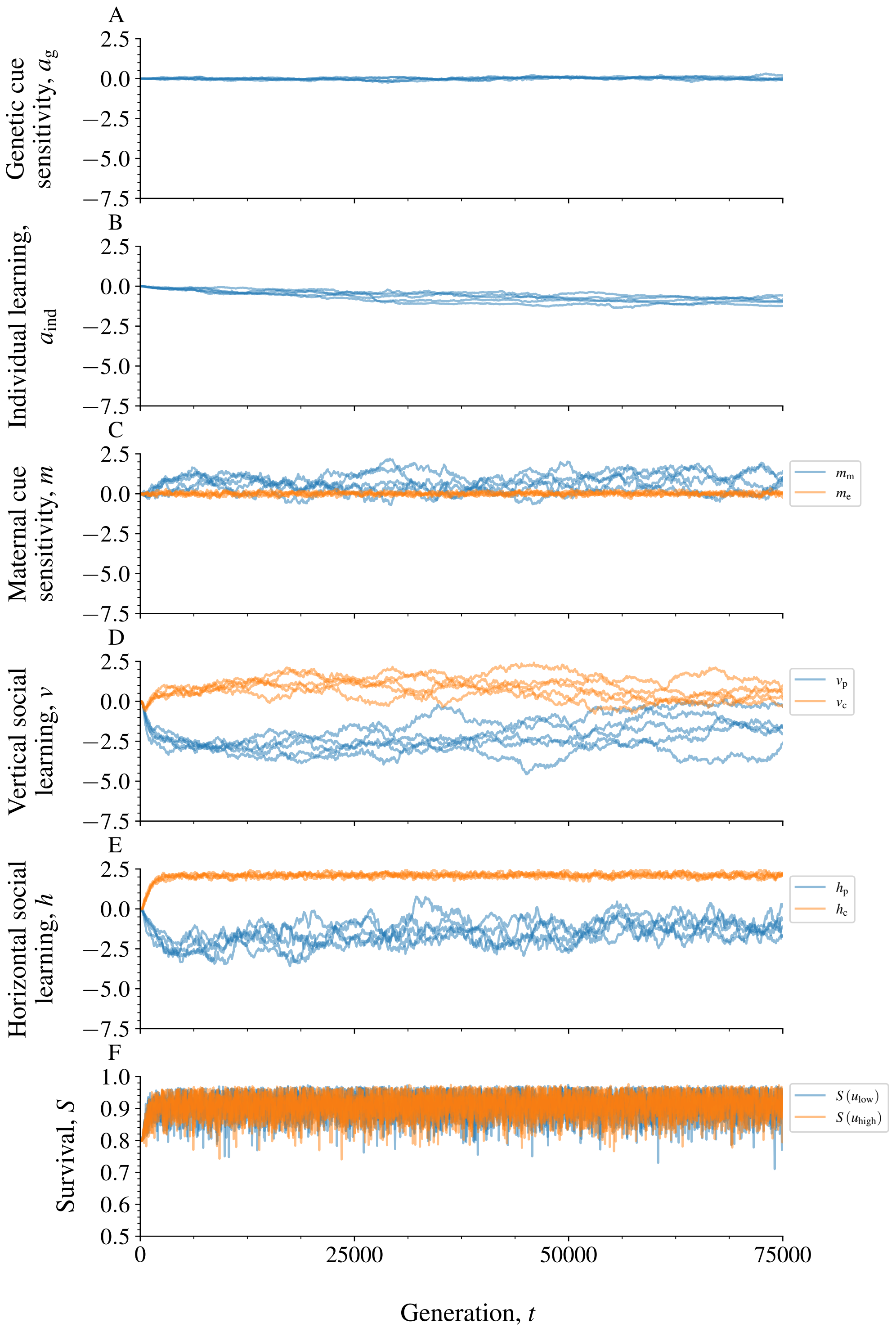
A
Genetic cue sensitivity, $a_{\mathrm{g}}$
B
Individual learning, $a_{\mathrm{ind}}$
C
Maternal cue sensitivity, $m$
$m_{\mathrm{m}}$
$m_{\mathrm{e}}$
D
Vertical social learning, $v$
$v_{\mathrm{p}}$
$v_{\mathrm{c}}$
E
Horizontal social learning, $h$
$h_{\mathrm{p}}$
$h_{\mathrm{c}}$
F
Survival, $S$
$S(u_{\mathrm{low}})$
$S(u_{\mathrm{high}})$
Generation, $t$

Figure S1:

**Figure S2** The evolved values of the sensitivities when varying the probability of environmental change $1-p$. See Figure 2 for the corresponding proportions of variance explained by the various sensitivity $\times$ cue combinations. See Figure 2 for parameters.

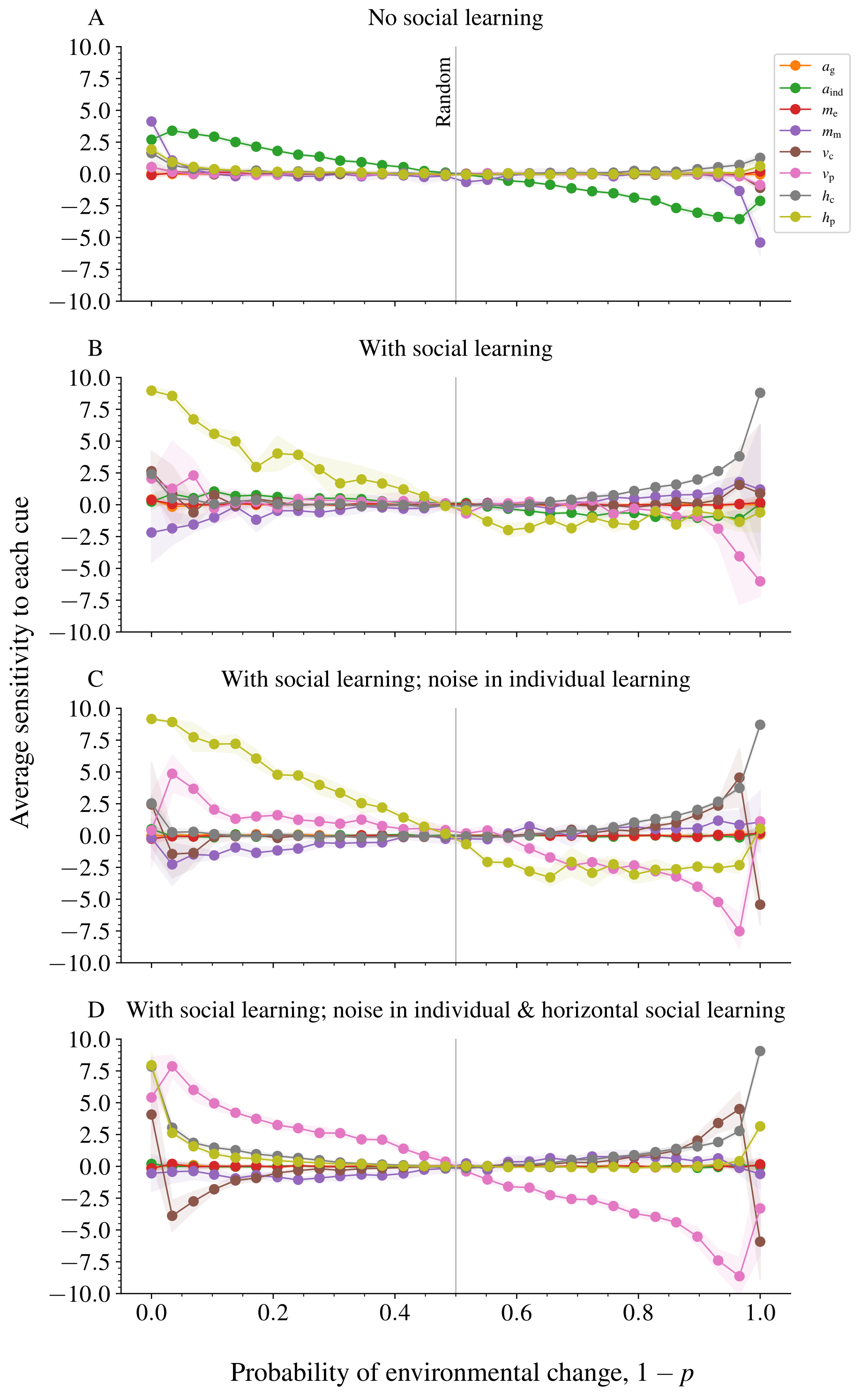
A
No social learning
Random
B
With social learning
C
With social learning; noise in individual learning
D
With social learning; noise in individual & horizontal social learning
10.0
7.5
5.0
2.5
0.0
−2.5
−5.0
−7.5
−10.0
0.0
0.2
0.4
0.6
0.8
1.0
Average sensitivity to each cue
Probability of environmental change, 1 − p
$a_g$
$a_{ind}$
$m_e$
$m_m$
$v_c$
$v_p$
$h_c$
$h_p$

Figure S2:

**Figure S3** Proportions of variance explained when horizontal social learning based on prestige or conformism biases coevolves exclusively with genetic cues (panel A), exclusively with maternal phenotypic cues (panel B) and without any other cues (panel C). See Figure 2 for parameters.

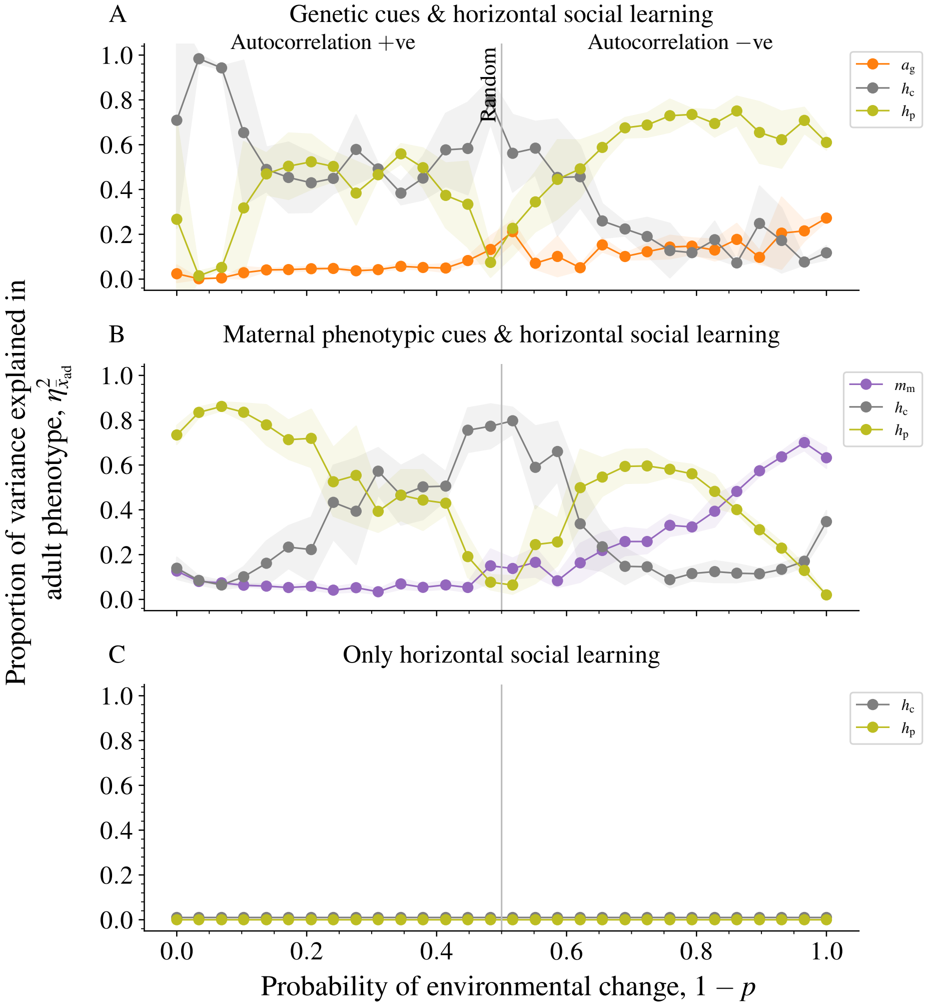
A
Genetic cues & horizontal social learning
Autocorrelation +ve
Autocorrelation −ve
Random
B
Maternal phenotypic cues & horizontal social learning
C
Only horizontal social learning
Proportion of variance explained in adult phenotype, $\eta^2_{\bar{x}_{ad}}$
Probability of environmental change, $1 - p$
$a_g$
$h_c$
$h_p$
$m_m$


Figure S3:

**Figure S4** Most important cues affecting juvenile phenotype determination (i.e., before horizontal social learning). Vertical prestige biases and individual learning mostly predominate. However, when fidelities of both vertical social learning and individual learning are low (top left corner of each panel), maternal effects typically prevail. Cases where vertical social learning based on conformity evolve (green areas) despite high levels of noise in vertical social learning merely reflect the generation of phenotypic variance: horizontal prestige-based learning (shown in Figure 3) evolves to strong values here, so that other cues are largely superfluous. See Figure 3 for parameters.

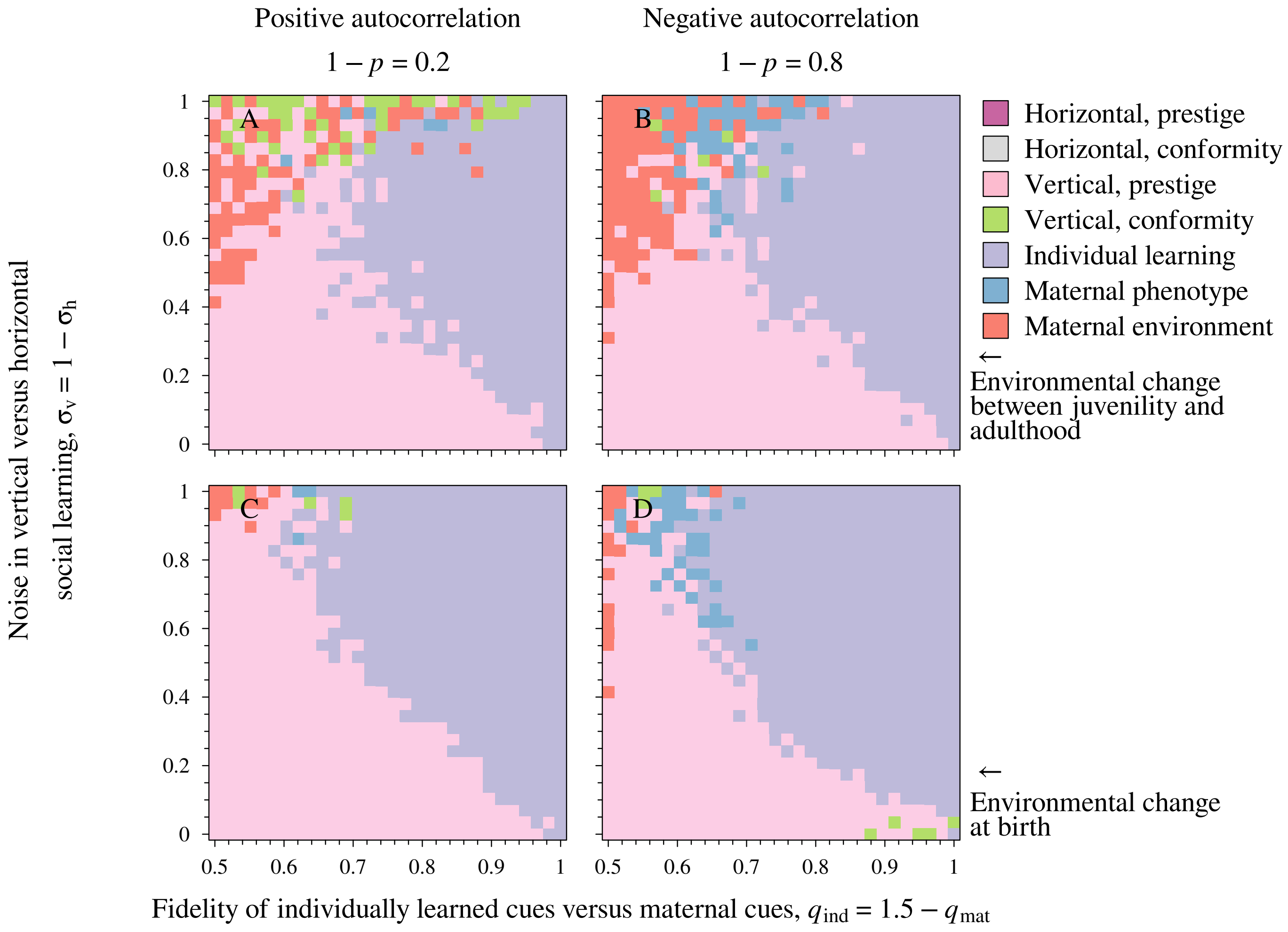
Positive autocorrelation
$1 - p = 0.2$
Negative autocorrelation
$1 - p = 0.8$
A
B
C
D
Horizontal, prestige
Horizontal, conformity
Vertical, prestige
Vertical, conformity
Individual learning
Maternal phenotype
Maternal environment
← Environmental change between juvenility and adulthood
← Environmental change at birth
Noise in vertical versus horizontal social learning, $\sigma_v = 1 - \sigma_h$
Fidelity of individually learned cues versus maternal cues, $q_{ind} = 1.5 - q_{mat}$
0
0.2
0.4
0.6
0.8
1
0.5
0.6
0.7
0.8
0.9
1

Figure S4:

**Figure S5** Proportions of variance explained when environmental change occurs at birth, rather than between juvenility and adulthood (compare with Figure 2). See Figure 2 for parameters.

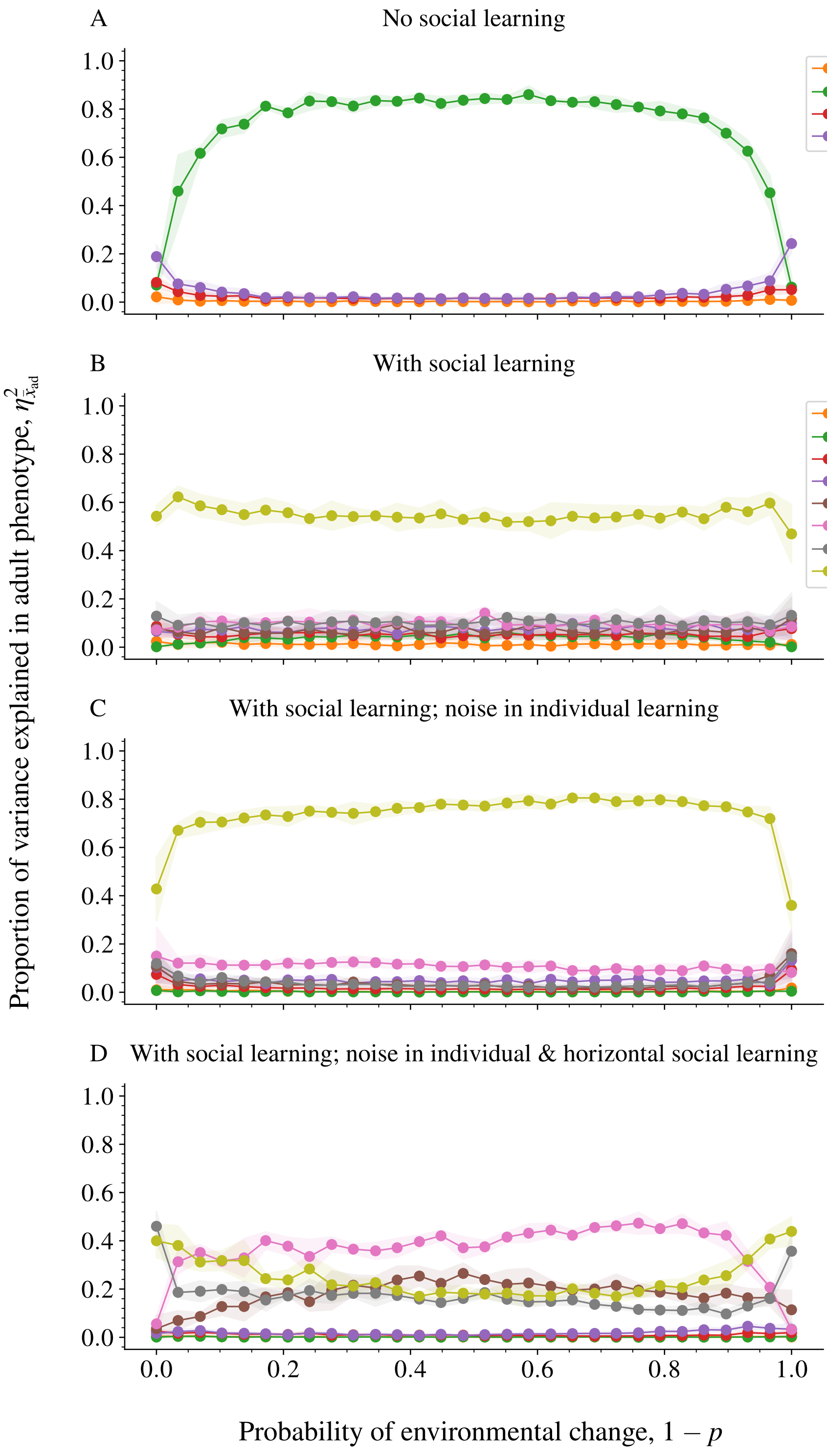
A
No social learning
B
With social learning
C
With social learning; noise in individual learning
D
With social learning; noise in individual & horizontal social learning
Proportion of variance explained in adult phenotype, $\eta^2_{\bar{x}_{ad}}$
Probability of environmental change, $1-p$
$a_g$
$a_{ind}$
$m_e$
$m_m$
$v_c$
$v_p$
$h_c$
$h_p$
0.0
0.2
0.4
0.6
0.8
1.0

Figure S5:

**Figure S6** Sensitivity analysis of the timing of vertical social learning and different values of migration $m$ (from left to right). The top two rows (panels A - F) consider the scenario discussed in the main text, in which juveniles learn vertically from breeders in the natal patch (i.e., vertical learning *before* migration), while they learn horizontally from other juveniles on the patch to which they have migrated (i.e., social learning *after* migration. By contrast, the bottom two rows (panels G - L) consider a scenario where both vertical and social learning occur from individuals on the patch to which a focal individual has migrated (i.e., learning *after* migration). Coloration again depicts the cue which explains the majority of phenotypic variance in adult phenotype (measured at the logistic scale).

When individuals learn vertically in their natal patch and learn horizontally in their patch of arrival (panels A - F), we find that for high levels of dispersal horizontal social learning prevails in both negatively and positively autocorrelated environments (panels C, F). This occurs because only horizontal learning allows an individual to obtain information about its current local environment.

By contrast, when individuals learn vertically as well as horizontally on their patch of arrival (panels G - L), horizontal social learning only prevails when it is characterised by low levels of noise, while vertical social learning prevails otherwise (compare panels I, L with C, F). In this case, both horizontal and vertical social learning (based on prestige) provide accurate information about the environment on the patch of arrival, so that prevalence of either is determined by the relative amount of noise $\sigma_v = 1 - \sigma_h$ that is varied on the $y$ axis. Other parameters as in Figure 3.

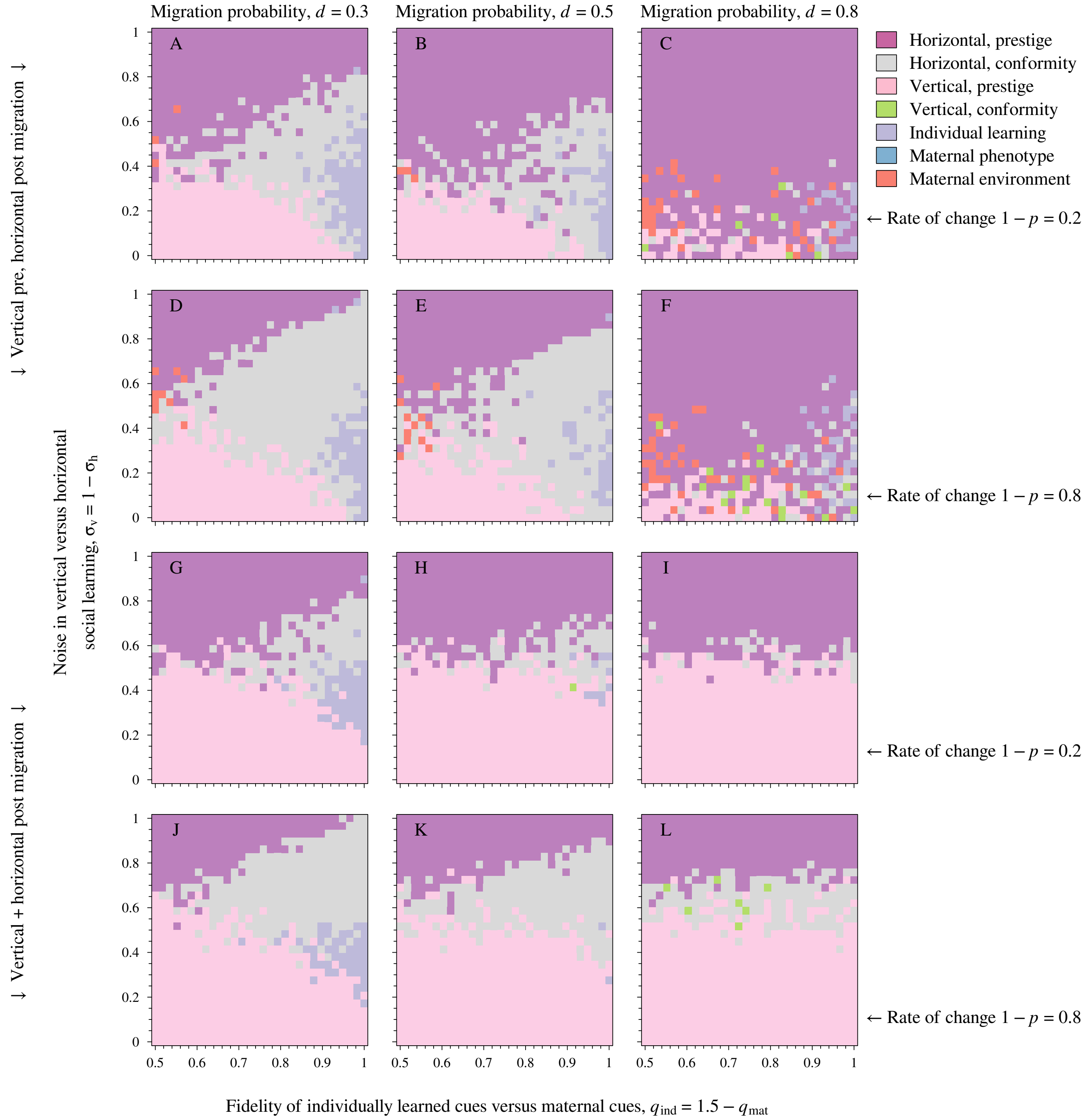

Migration probability, $d = 0.3$
Migration probability, $d = 0.5$
Migration probability, $d = 0.8$
A
B
C
D
E
F
G
H
I
J
K
L
Horizontal, prestige
Horizontal, conformity
Vertical, prestige
Vertical, conformity
Individual learning
Maternal phenotype
Maternal environment
← Rate of change $1 - p = 0.2$
← Rate of change $1 - p = 0.8$
← Rate of change $1 - p = 0.2$
← Rate of change $1 - p = 0.8$
↓ Vertical pre, horizontal post migration ↓
↓ Vertical + horizontal post migration ↓
Noise in vertical versus horizontal social learning, $\sigma_v = 1 - \sigma_h$
Fidelity of individually learned cues versus maternal cues, $q_{ind} = 1.5 - q_{mat}$
0
0.2
0.4
0.6
0.8
1
0.5
0.6
0.7
0.8
0.9
1


Figure S6:

**Figure S7** Sensitivity analysis of the timing of horizontal learning (panels A - F), the timing of individual learning (panels G - L) and different values of migration $m$ (from left to right).

The top two rows (panels A - F) consider a scenario in which individuals perform all their learning before migration: individual learning and vertical social learning is assumed to occur before migration as in the main text, while now also horizontal social learning happens before migration As a consequence, individuals only acquire information about the environmental state of their natal patch. With an increasing probability of migration, individuals thus lack any information about their future environment, resulting in an outcome where all cues are equally informative (panels C, F) as testified by the large variation between individual simulations in the cue that is most informative.

The bottom two rows (panels G - L) consider the same scenario as in the main text (vertical social learning prior to migration, horizontal social learning after migration), except that individual learning now occurs after dispersal. As a consequence, individual learning (once it has a high fidelity as measured by $q_{\text{ind}}$ on the left-hand side of each panel) now prevails over a much larger part of the parameter space, particularly when dispersal is high (i.e., compare panels I, L with Figure S6C, F). Other parameters as in Figure 3.

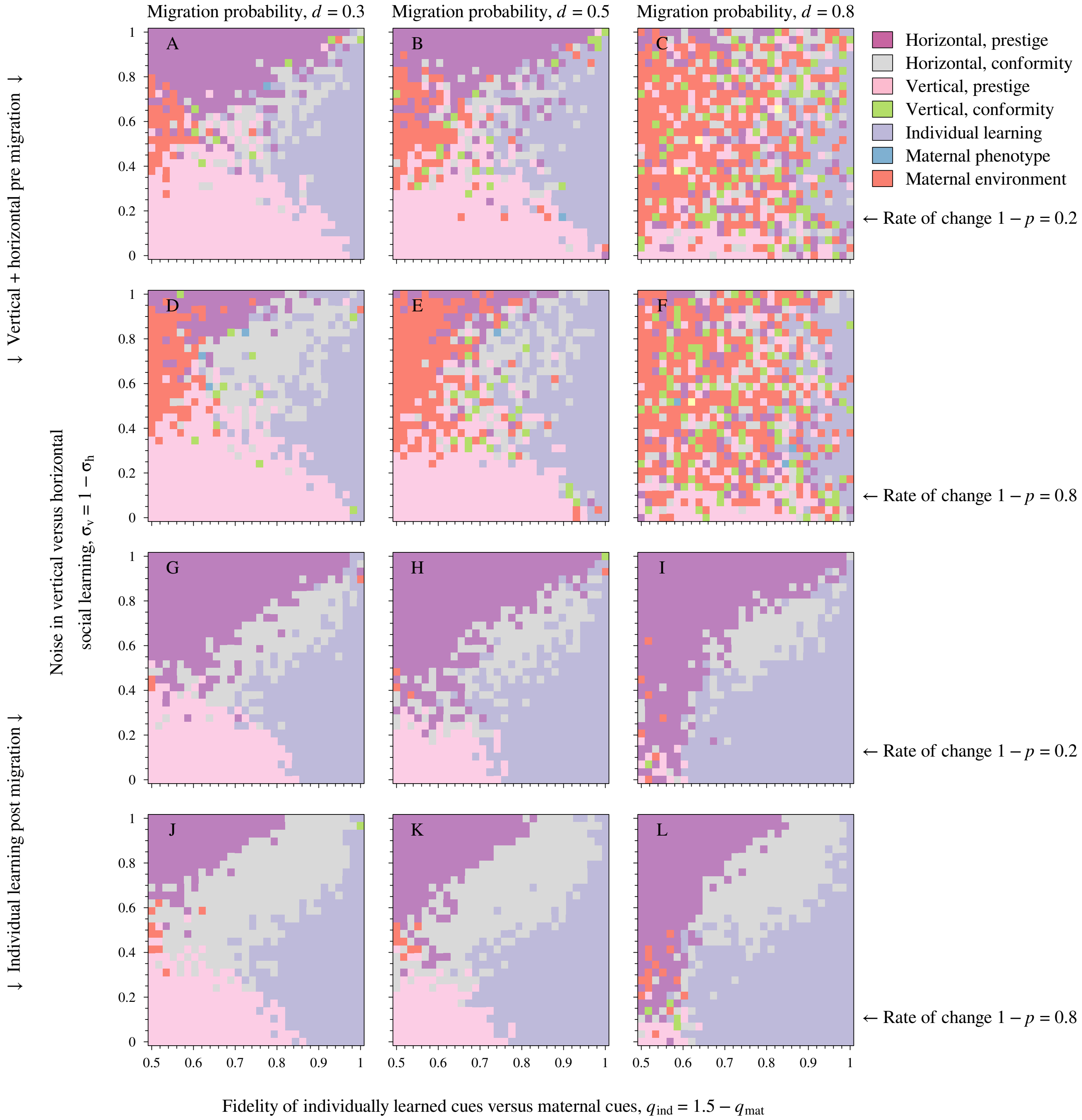
Migration probability, $d$ = 0.3
Migration probability, $d$ = 0.5
Migration probability, $d$ = 0.8
Horizontal, prestige
Horizontal, conformity
Vertical, prestige
Vertical, conformity
Individual learning
Maternal phenotype
Maternal environment
← Rate of change $1 - p = 0.2$
← Rate of change $1 - p = 0.8$
← Rate of change $1 - p = 0.2$
← Rate of change $1 - p = 0.8$
↓ Vertical + horizontal pre migration ↓
↓ Individual learning post migration ↓
Noise in vertical versus horizontal social learning, $\sigma_v = 1 - \sigma_h$
Fidelity of individually learned cues versus maternal cues, $q_{ind} = 1.5 - q_{mat}$
A
B
C
D
E
F
G
H
I
J
K
L

Figure S7:

**Figure S8** The effect of varying the number $n$ of models sampled when performing horizontal or vertical social learning. As in Figure 3 in the main text (in which $n = 5$). Panel A: when $n = 1$ (i.e., random sampling of models), we find that maternal environmental effects or individual learning prevail, while conformity biases prevail at intermediate values of $q_{\text{ind}} = 1.5 - q_{\text{mat}}$, effectively reflecting social learning from random individuals. Once $n$ gets larger, we find that maternal effects and individual learning are replaced by prestige-based social learning (either vertical or horizontal). Parameters as in Figure 3.

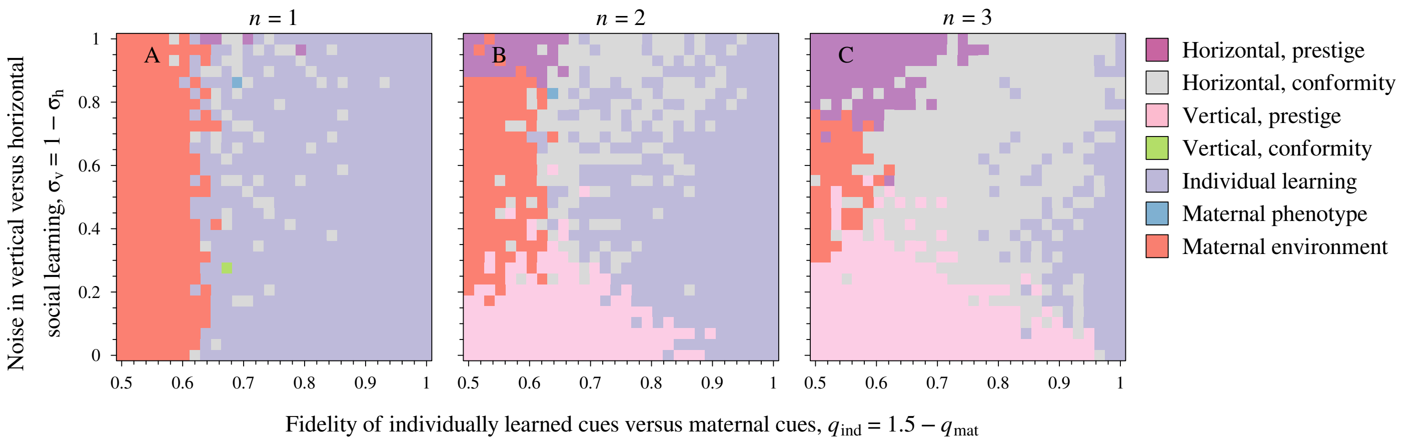


Figure S8:

**Figure S9** Cues that explain the majority of phenotypic variance in adult phenotype (see Figure 3 in the main text) when social learning is only based on conformity biases. In comparison to Figure 3, we now find that horizontal prestige-based social learning is replaced by horizontal learning based on conformity biases. By contrast, vertical prestige-based social learning is replaced either by maternal effects (towards left-hand sign of each panel) or individual learning once its fidelity is large enough. Parameters as in Figure 3.

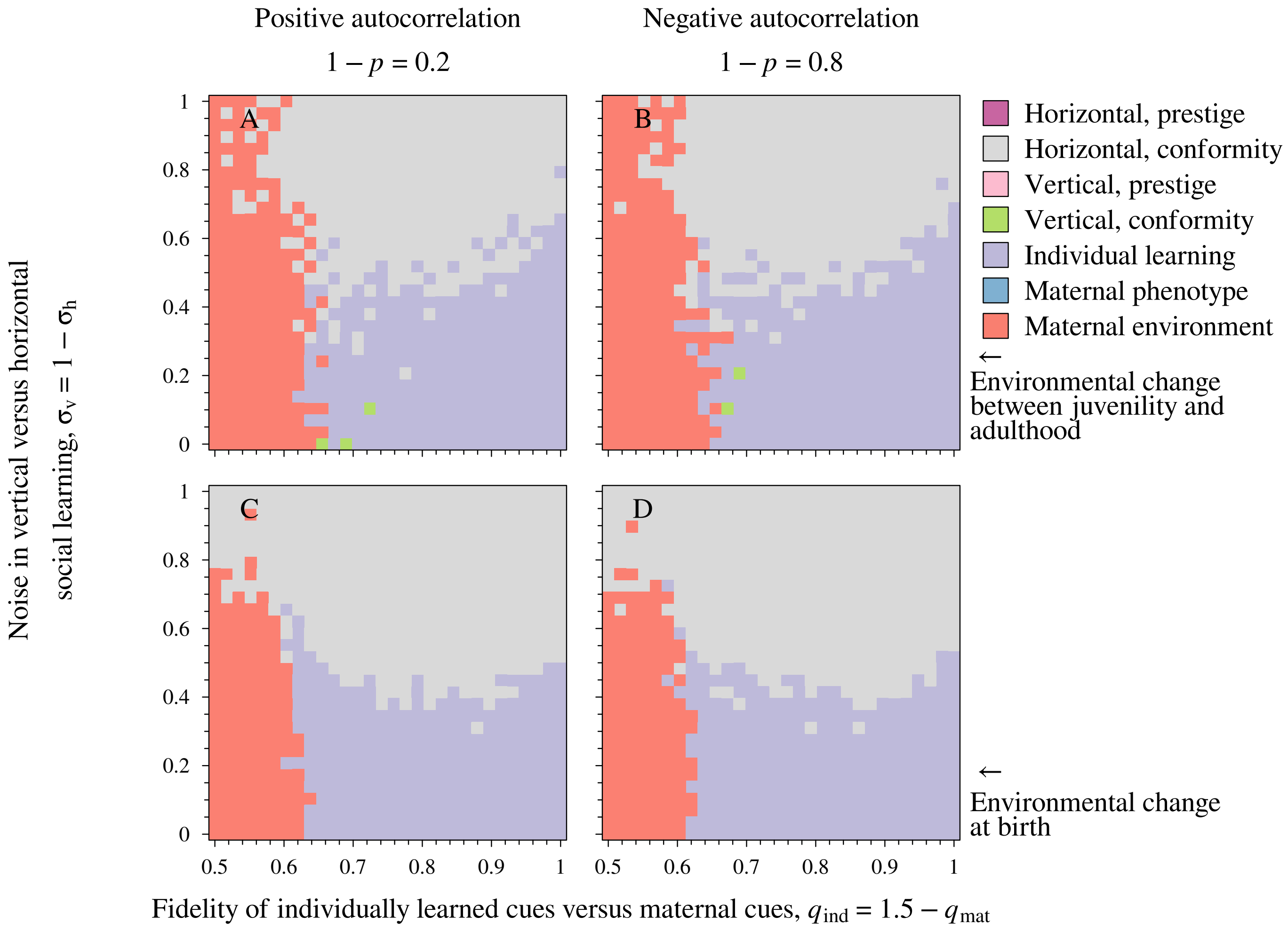
Positive autocorrelation
1 − p = 0.2
Negative autocorrelation
1 − p = 0.8
A
B
C
D
Horizontal, prestige
Horizontal, conformity
Vertical, prestige
Vertical, conformity
Individual learning
Maternal phenotype
Maternal environment
← Environmental change between juvenility and adulthood
← Environmental change at birth
Noise in vertical versus horizontal social learning, σv = 1 − σh
Fidelity of individually learned cues versus maternal cues, qind = 1.5 − qmat
0
0.2
0.4
0.6
0.8
1
0.5
0.6
0.7
0.8
0.9
1

Figure S9:

**Figure S10** Here we assess the impact of environmental frequencies that are different from $1/2$ on the evolution of cue integration. The figure varies (i) the frequency of the two environments, (ii) the autocorrelation and (iii) the fidelity of individually learned cues (panels A vs B) for the sake of comparison with Figure 2 in the main text. For autocorrelations at or near 0, we find that neither cue is informative (conservative bet-hedging) so that any cue can prevail (see also Figure 2 when $1-p=0.5$). By contrast when autocorrelations are positive, we find that horizontal learning based on prestige biases prevails as on the left-hand side in Figure 2B, C. When autocorrelations are negative, we find that vertical social learning typically prevails (in combination with horizontal conformity-based social learning) when the fidelity of individual learning is low (panel A, similar to the right-hand side of Figure 2C), while individual learning prevails otherwise (panel B, similar to the right-hand side of Figure 2B). Hence, these results are very similar to what has been found in Figure 2. In positively autocorrelated environments, horizontal learning based on prestige biases prevails, again as in Figure 2.

To generate this figure, we have replaced the probability of environmental change $1-p$ used in the main text by environment-specific probabilities of environmental change, so that for an individual patch in environments 1 and 2 the per-generation probabilities of change to a different environmental state are given by $\mathcal{S}_{1\to 2}$ and $\mathcal{S}_{2\to 1}$ respectively. Consequently, the expected frequency of environment 1 is given by $\mathcal{S}_{2\to 1}/(\mathcal{S}_{1\to 2}+\mathcal{S}_{2\to 1})$ while the autocorrelation of any local environment between the current generation and the next is given by $1-(\mathcal{S}_{1\to 2}+\mathcal{S}_{2\to 1})$. Combinations in which there is a low frequency of either environment, while a strongly negative autocorrelation are not feasible as they result in values of $\mathcal{S}_{i\to j}$ that are outside of the $[0,1]$ range, as indicated by white areas.

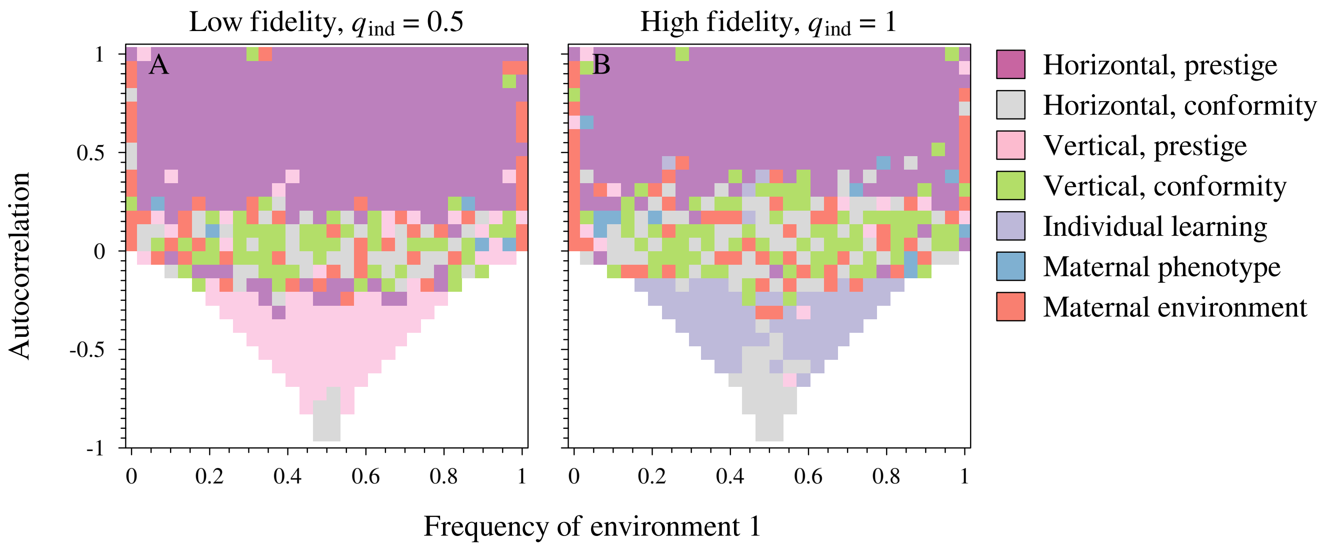

Low fidelity, $q_{\text{ind}} = 0.5$
High fidelity, $q_{\text{ind}} = 1$
A
B
1
0.5
0
-0.5
-1
Autocorrelation
0
0.2
0.4
0.6
0.8
1
Frequency of environment 1
Horizontal, prestige
Horizontal, conformity
Vertical, prestige
Vertical, conformity
Individual learning
Maternal phenotype
Maternal environment


Figure S10:

### S2 Supplementary model description

#### S2.1 Survival selection

We assume that survival selection $S(u,\theta_i)$ of adults is given by the function (see also eqns. [1,2] in [17]):

$$S(u,\theta_i) = \begin{cases} 1-0.8u^2 & \theta_i = \theta_{\text{low}} \\ 1-0.8(1-u)^2 & \theta_i = \theta_{\text{high}} \end{cases}, \tag{S1}$$

which is the function depicted in Figure 1B.

#### S2.2 Juvenile phenotype determination

To recap from the main text, the juvenile phenotype $u_{\text{juv}}$ that is developed after individual learning is a logistic function of a weighted sum $\bar{x}_{\text{juv}}$ of different cues an individual has received. We have

$$u_{\text{juv}} = \frac{1}{1+\exp\left(-\bar{x}_{\text{juv}}\right)} \tag{S2}$$

$$\begin{aligned} \bar{x}_{\text{juv}} = {} & a_{\text{g}}x_{\text{gen}} + a_{\text{ind}}x_{\text{ind}} \\ & + m_{\text{m}}x_{\text{mat,phen}} + m_{\text{e}}x_{\text{mat,envt}} \\ & + v_{\text{p}}x_{\text{vert,prestige}} + v_{c}x_{\text{vert,conformity}}, \end{aligned} \tag{S3}$$

where the $x_i$s in the equation above are the values of each of the different cues, each of which is weighed by a sensitivity locus that can flexibly evolve. Regarding the evolving sensitivity loci, $a_{\text{g}}$ is the sensitivity to the genetic cue $x_{\text{gen}}$ (see section S2.2.1 below). The locus $a_{\text{ind}}$ reflects evolving sensitivity to individually learned cues $x_{\text{ind}}$ that inform about the state of local environment at birth (see section S2.2.2 below). The locus $m_{\text{m}}$ reflects evolving sensitivity to the maternal phenotype $x_{\text{mat,phen}}$ as a cue, while the locus $m_{\text{e}}$ reflects evolving sensitivity to the maternal environment $x_{\text{mat,envt}}$ as a cue (see section S2.2.3). Finally, the juvenile phenotype is also influenced by vertical social learning: here, $v_{\mathrm{p}}$ reflects evolving sensitivity to vertically learned phenotypic cues based on prestige biases $x_{\mathrm{vert,prestige}}$ (see section S2.2.4). The locus $v_{\mathrm{c}}$ reflects evolving sensitivity to vertically learned phenotypic cues based on conformity biases (see section S2.2.4).

##### S2.2.1 Genetic cues

Following similar models by [12, 15, 17], the value of the genetic cue $x_{\mathrm{gen}}$ is given by $x_{\mathrm{gen}} = \sum_{i=0}^{n_{\mathrm{g}}} g_i$. That is, it is the sum of the allelic values of $n_{\mathrm{g}} = 3$ unlinked, diploid genetic cue loci $g_i$. Each allele can have values in the range of $-1 \leq g_i \leq 1$. Consequently, limited dispersal can result in scenarios where alleles become associated with the local environment, so that alleles become informative. Note that alleles of the genetic cue themselves are not involved in local adaptation, they purely have an informational function. The role of genetic cues is further discussed in [13].

##### S2.2.2 Individually learned cues

Juveniles perform individual learning about the state of their local environments by observing a juvenile environmental cue $x_{\mathrm{ind}}$. To this end, each individual independently observes a binary cue reflecting the state of the local environment. The cue is equal to the actual environmental state (value $-1/2$ in environment $\theta_{\mathrm{low}}$, value $1/2$ in environment $\theta_{\mathrm{high}}$) with a probability given by the cue fidelity parameter $0.5 \leq q_{\mathrm{juv}} \leq 1$, whereas with probability $1-q_{\mathrm{juv}}$ an individual receives a cue associated with the opposite environmental state.

##### S2.2.3 Maternal cues

Juveniles receive two maternal cues: the first is a maternal phenotypic cue $x_{\mathrm{mat,phen}} = u_{\mathrm{ad}}(t-1)-1/2$ that reflects the value of the maternal phenotype $0 \leq u_{\mathrm{ad}}(t-1) \leq 1$. By subtracting $1/2$ from $u_{\mathrm{ad}}(t-1)$ we standardise the maternal phenotype between $-1/2$ and $1/2$ so that the maternal phenotypic cue has the same range as other plastic cues (and hence evolved cue sensitivities can be directly compared). As $u_{\mathrm{ad}}(t-1)$ is itself a function of her mother's phenotype $u_{\mathrm{ad}}(t-2)$ (a so-called cascading parental effect: [16, 46, 47]), a maternal phenotypic cue can potentially give rise to grandmaternal effects. See [17] for an assessment of noise in maternal phenotypic cues, which we do not consider here for the sake of brevity.

The second maternal cue is a maternal environmental cue $x_{\text{mat,envt}}$, which reflects the value of the maternal environment [48] as provided to the mother by the adult environmental cue (see Figure 1). The cue is equal to the actual environmental state (value $-1/2$ in environment $\theta_{\text{low}}$, value $1/2$ in environment $\theta_{\text{high}}$) with a probability given by the cue fidelity parameter $0.5 \leq q_{\text{mat}} \leq 1$, whereas with probability $1 - q_{\text{mat}}$ an individual receives a cue associated with the opposite environmental state. We assume that all offspring born form the same mother obtain the same value of $x_{\text{mat,envt}}$ (i.e., errors act at the level of the brood, rather than at the level of the individual offspring).

###### S2.2.4 Vertical social learning: prestige biases

Following previous models of prestige-based learning [e.g., 49, 50], we assume that individuals are able to evaluate and rank the performance of observed models. To this end, learners rank the potential survival payoffs $S(u_{\text{ad},i})$ of each individual $i$ in a random sample of $n_{\text{p}}$ adult models after survival selection has taken place in the previous generation (see Figure 1). The potential survival payoff is evaluated based on $i$th model's adult phenotype $u_{\text{ad},i}$ that is observed by the learner (see eq. [3] below) in the learner's environment at time of birth (see Figure 1). The cue $x_{\text{vert,prestige}}$ is then a function of the phenotype $u_{\text{ad,max}}$ of the sampled individual which has the highest ranked survivorship value, namely $x_{\text{vert,prestige}} = u_{\text{ad,max}} + \xi_{\text{vert,prestige}} - 1/2$. Here $\xi_{\text{vert,prestige}}$ is a sample drawn from a Gaussian noise distribution with mean 0 and standard deviation $\sigma_{\text{vert,prestige}}$, and subtracting $-1/2$ standardises the cue range to the same scale as for other cues.

###### S2.2.5 Vertical social learning: conformity-biases

We follow previous models of conformity-based social learning [e.g., 41] where individuals evaluate phenotypes $u_i$ of each individual belonging to randomly chosen subset of $n_{\text{c}}$ surviving adult models from the parental generation in the local patch. Individuals take account of the number of individuals $n_{\text{c,lo}} \leq n_{\text{c}}$ with a phenotypes $u_i$ corresponding to the low environment (i.e., $u_i < 0.5$), th

Figure S11:

whereas $n_{\text{c,hi}} = n_{\text{c}} - n_{\text{c,lo}}$ reflects the number of individuals with phenotypes $u_i \geq 0.5$. We then have

$$x_{\text{vert,conformity}} = \begin{cases} -1/2 & n_{\text{c,lo}} > n_{\text{c,hi}} \text{ (low-matching phenotype predominates)} \\ 0 & n_{\text{c,lo}} = n_{\text{c,hi}} \text{ (no phenotype predominates)} \\ 1/2 & n_{\text{c,lo}} < n_{\text{c,hi}} \text{ (high-matching phenotype predominates)} \end{cases} .$$

Moreover, we add noise to the cue by adding a random deviate from a Gaussian distribution with mean 0 and standard deviation $\sigma_{\text{vert,conformity}}$. We have also considered alternative configurations where $x_{\text{vert,conformity}} = n_{\text{c,hi}}/(n_{\text{c,hi}} + n_{\text{c,lo}})$, but these give similar results (results not shown).

#### S2.3 Adult phenotype determination

To recap from the main text, we have

$$u_{\text{ad}} = \frac{1}{1+\exp(-\bar{x}_{\text{ad}})} \tag{S4}$$

$$\bar{x}_{\text{ad}} = \bar{x}_{\text{juv}} + h_{\text{p}} x_{\text{horiz,prestige}} + h_c x_{\text{horiz,conformity}}, \tag{S5}$$

where $h_{\text{p}}$ (horizontal social learning; prestige bias) and $h_{\text{c}}$ (horizontal social learning; conformity bias) again reflect unlinked and evolving diploid loci (bounded between [-10,10]) that reflect sensitivity to both horizontally learned social cues $x_{\text{horiz,prestige}}$ and $x_{\text{horiz,conformity}}$ respectively.

##### S2.3.1 Horizontal social learning: prestige biases

Horizontally learning juveniles rely on the same mechanism as vertical social learning, where they rank the potential survival payoffs $S(u_{\text{juv},i})$ of each individual $i$ in a random sample of $n_{\text{p}}$ juvenile models (no survival selection has yet taken place; see Figure 1). The potential survival payoff is evaluated based on $i$ model's juvenile phenotype $u_{\text{juv},i}$ (see eq. S2) that is observed by the juvenile learner in the learner's environment at time of birth (see Figure 1). The cue $x_{\text{horiz,prestige}}$ is then a function of the phenotype $u_{\text{juv,max}}$ of the sampled individual which has the highest ranked survivorship value, namely $x_{\text{horiz,prestige}} = u_{\text{juv,max}} + \xi_{\text{horiz,prestige}} - 1/2$. Here $\xi_{\text{horiz,prestige}}$ is a sample drawn from a Gaussian noise distribution with mean 0 and standard deviation $\sigma_{\text{horiz,prestige}}$, and subtracting $-1/2$ standardises the cue range to the same scale as for other cues.

##### S2.3.2 Horizontal social learning: conformity-biases

We follow previous models of conformity-based social learning [e.g., 41] where individuals evaluate phenotypes $u_i$ of each individual belonging to randomly chosen subset of $n_{\text{c}}$ juvenile models from the current generation in the local patch. Individuals take account of the number of individuals $n_{\text{c,lo}} \leq n_{\text{c}}$ with a phenotypes $u_i$ corresponding to the low environment (i.e., $u_i < 0.5$), whereas $n_{\text{c,hi}} = n_{\text{c}} - n_{\text{c,lo}}$ reflects the number of individuals with phenotypes $u_i > 0.5$. We then have

$$x_{\text{horiz,conformity}} = \begin{cases} -1/2 & n_{\text{c,lo}} > n_{\text{c,hi}} \text{ (low-matching phenotype predominates)} \\ 0 & n_{\text{c,lo}} = n_{\text{c,hi}} \text{ (no phenotype predominates)} \\ 1/2 & n_{\text{c,lo}} < n_{\text{c,hi}} \text{ (high-matching phenotype predominates)} \end{cases}.$$

Moreover, we add noise to the cue by adding a random deviate from a Gaussian distribution with mean 0 and standard deviation $\sigma_{\text{horiz,conformity}}$.

#### S2.4 Inheritance

For the sake of simplicity, the gene loci coding for the cue sensitivities $a_i$, $m_i$, $h_i$ and $v_i$ and the genetic cue loci $g_i$ are all considered diploid, autosomal and unlinked. Upon inheritance, alleles at all loci independently mutate with probability $\mu = 0.01$, which involves adding a random value drawn from a Laplace$(\mu, b)$ distribution with parameters $\mu = 0$ and $b = \sigma_\mu/\sqrt{2} = 0.0141$, corresponding to a mean of 0 and a variance of $\sigma_\mu^2 = 0.0004$. To ensure the genetic cue locus can accumulate sufficient genetic variation in, we assume an increased mutational variance $\sigma_{\mu_g}^2 = 0.0625$ for the alleles at the genetic cue locus $g_i$.
